# SCORCH2: a generalised heterogeneous consensus model for high-enrichment interaction-based virtual screening

**DOI:** 10.1101/2025.03.31.646332

**Authors:** Lin Chen, Vincent Blay, Pedro J. Ballester, Douglas R. Houston

## Abstract

The discovery of effective therapeutics remains a complex, costly, and time-consuming endeavor, characterized by high failure rates and significant resource investments. A central bottleneck in early-stage drug discovery is identifying suitable hit compounds with moderate affinity for known biological targets. Although advancements have been made, current *in silico* virtual screening methods are subject to limitations, including model overfitting, data bias, and constrained interpretability in their predictive processes. In this study, we present SCORCH2, a machine learning-based framework designed to enhance both the performance and interpretability of virtual screening by leveraging interaction features. Compared with its predecessor SCORCH, SCORCH2 exhibits superior predictive accuracy and generalizability across a wide range of biological targets. Importantly, SCORCH2 demonstrates robust hit identification capabilities on previously unseen targets, indicating strong transferability. These results highlight the potential of SCORCH2 as a valuable tool in accelerating drug discovery, offering reliable predictive capabilities while improving the interpretability of virtual screening models.

## 1 Main

The rising demand for effective treatments underscores the importance of identifying novel therapeutic agents. A key determinant of a compound’s therapeutic potential is its binding affinity, which can be experimentally quantified using metrics such as the dissociation constant (*K*_*d*_), inhibition constant (*K*_*i*_), or Gibbs free energy change (Δ*G*) [1]. However, identifying viable drug candidates without experimental validation remains a formidable challenge. In particular, early-stage drug discovery is often hindered by the difficulty of discovering initial “hit” compounds that exhibit the desired biological activity against a predefined target. This task is further complicated by the vast and sparsely populated nature of chemical space [2, 3], which limits the efficiency of exhaustive experimental approaches. To address these limitations, virtual screening (VS) techniques have emerged as computational alternatives to traditional high-throughput screening. These methods aim to reduce the time and cost associated with drug discovery by leveraging large datasets and machine learning algorithms to improve both the efficiency and accuracy of hit identification [4–9]. Despite significant progress, the reliability and general applicability of current *in silico* models remain constrained by several persistent challenges. These include:

1. Unrealistic performance [10]. Existing benchmarks such as DUD-E [11] have been shown to introduce substantial dataset biases, resulting in inflated performance metrics that fail to generalize to more diverse datasets such as DEKOIS [12, 13]. These biases obscure the true predictive capacity of virtual screening models [10].
2. Reliance on structural similarity: Model performance can be unduly influenced by protein sequence or ligand similarity, leading to a phenomenon of memorization rather than true predictive power[14–16].
3. Uncertainty in evaluation: Although training data incorporating experimentally validated crystal structures can improve pose quality, many widely used benchmarks (DUD-E, DEKOIS, LIT-PCBA[17]) lack crystal pose information for validation data. Retrospective analyses of the PDBBind coreset indicate that existing docking methods do not reliably generate near-native docking poses (RMSD ≤ 2 Å), nor do they consistently rank the most near-native pose as the top prediction[18–21]. This introduces a potential for inaccurate predictions stemming from errors in the objective docking poses.
4. Annotation scarcity: The financial and logistical constraints associated with the experimental determination of binding affinity restrict the availability of annotated data, thereby limiting its utility in practical applications.

A promising approach involves reframing virtual screening (VS) as a specialized scoring function (SF) application, focusing on the assessment of ligand-receptor binding mode plausibility rather than the prediction of general alchemical properties. This strategy leverages experimentally determined complex structures, derived from techniques like X-ray crystallography and cryo-electron microscopy (Cryo-EM)[22], to characterize the energy landscape associated with productive binding, without requiring detailed experimental annotations. This perspective presents several benefits. First, it facilitates the broader use of unlabeled biological data, thereby addressing the limitations imposed by annotation scarcity. Second, it avoids data imbalance issues by assessing the ligand’s binding potential and treating compounds with varying potencies equitably. A key consideration, however, is the interpretability of the learned features. Empirical scoring functions like Vina [23] are commonly used, which rank ligands according to weighted combinations of energy terms. While attempts have been made to reparameterize these energy terms for VS, these efforts have often fallen short in capturing the complexities of structural and interaction data[24]. Furthermore, real-world VS applications often deal with highly imbalanced data distributions. To address this, model robustness can be improved by incorporating decoy molecules into the training set[25, 26]. These decoys, selected for their structural dissimilarity to known binders, are presumed not to bind the target. It is important to note that the use of decoys differs from methods focused on binding affinity prediction, as decoys do not provide information regarding binding strength. These distinctions emphasize the need to treat virtual screening as a distinct task from affinity prediction and highlight the need for new scoring methods that accurately represent the non-linear relationships between ligands and decoys.

To address the aforementioned challenges, we introduce SCORCH2 (SC2), the next-generation iteration of SCORCH[4](SC1). SC2 builds upon the feature set of SC1 by incorporating additional molecular descriptors and utilizing more diverse training datasets to capture a wider range of interaction patterns, thus enhancing its generalization capacity. SC2 has undergone rigorous evaluation using three established benchmarks—DUD-E, DEKOIS 2.0, and VSDS-vd by employing publicly available docking poses. The results suggest that SC2 effectively prioritizes highly-ranked candidates based primarily on ligand-protein interaction features and molecular descriptors, without dependence on explicit structural or sequence information. Comparative assessments demonstrate that SC2 achieves performance comparable to or exceeding that of current state-of-the-art (SOTA) methods, especially when applied to novel targets. Furthermore, SHAP (SHapley Additive exPlanations) feature importance analysis [27] indicates that SC2 prioritizes chemically relevant features, such as hydrogen bond formation and key energy terms, aligning its predictive behavior with established biochemical principles.

## 2 Results

### SCORCH2 overview

SC2 is a machine learning model designed for interaction-based virtual screening (IBVS). It is trained using binary labels to differentiate between active ligands and decoys by re-evaluating docking poses produced by external docking software. Consistent with other contemporary VS approaches, such as RTMScore[6] and EquiScore[5], SC2 seeks to establish a scoring framework where the predicted output represents a measure of binding confidence, as opposed to a quantitative prediction of binding affinity. Figure 1 depicts the SC2 framework and the associated workflow for both training and inference phases.

**Fig. 1.**
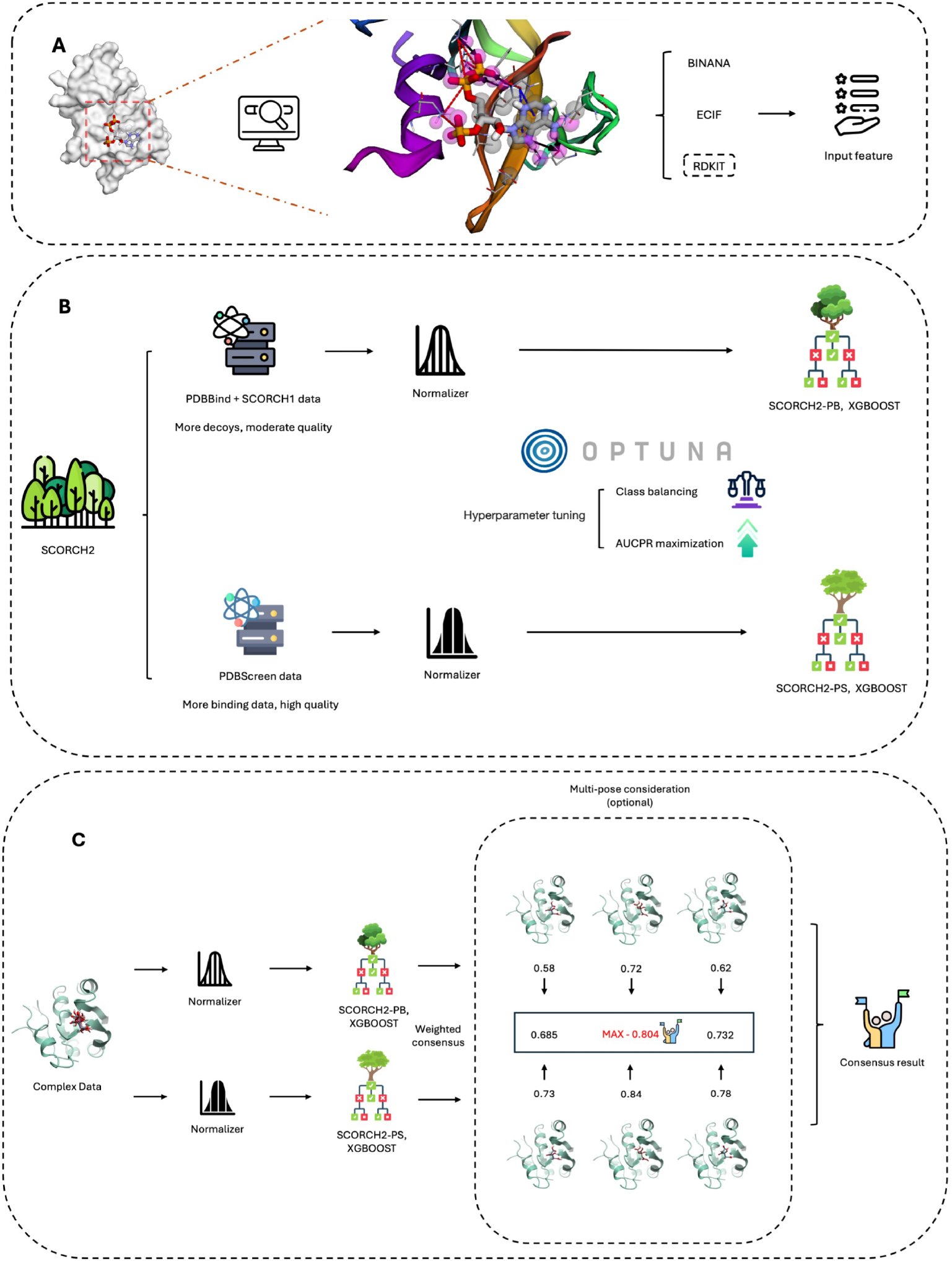
A, Molecular interaction visualization and feature combination. Crystal structure of protein with PDB ID: 1AFK, highlighting interactions with the ligand PAP (3’-Phosphate-Adenosine-5’-Diphosphate). Fuchsia spheres represent atoms in closest contact with the ligand (cutoff within 2.5 Å). Black arrows indicate hydrogen bonds, red dashed lines represent salt bridges, and blue dashed lines denote *π*–*π* stacking interactions. SCORCH2 features are majorly generated from three methods BINANA, ECIF, and RDKIT. B, SCORCH2 structure and model training. SC2 utilizes a simplified architecture consisting of two distinct XGBoost models. Each model is fed with different data, and Optuna is used for optimal parameter search with appropriate training strategies. C, SCORCH2 inference workflow, for the complex data, these models are designed to operate independently and the final result is provided by a weighted consensus.

SC2 is distinguished from prior VS models by three characteristics. First, through the utilization of distinct training datasets and data curation strategies, SC2 comprises two separate XGBoost[28] models for consensus scoring. These models capture divergent knowledge patterns, offering a more comprehensive assessment of binding potential from multiple perspectives. Each model can be employed independently, or their results can be aggregated to enhance generalisability. Second, SC2 represents a physics-inspired *in silico* IBVS model, a specific category within structure-based virtual screening (SBVS). Unlike conventional SBVS approaches that explicitly model the receptor (for instance, by constructing protein graphs in graph neural networks), SC2 treats the receptor as an implicit information medium. It leverages the three-dimensional structure solely for the extraction of ligand-receptor interaction features, and SC2 does not include protein descriptors. Finally, because of the primacy of molecular interactions in its predictive process, SC2 typically exhibit a lower number on AUROC (Area Under the Receiver Operating Characteristic curve) compared to analogous models; however, it achieves superior top-tier enrichment, highlighting its distinctive capability in identifying active compounds.

In the subsequent sections, we evaluate the performance of SC2 across various aspects. Given that SC2’s decisionmaking process relies heavily on non-covalent molecular interactions, it is inherently sensitive to pose quality; consequently, results may vary depending on the quality of the poses, the availability of compounds from the docking software, the pose sampling strategy employed, and other factors. To ensure standardized and equitable comparisons, we adopted the pose set utilized in EquiScore [5], where the docking poses are publicly accessible. Evaluations are performed using these retrieved docking poses; unless otherwise specified, reported SC2 results are based on Glide SP docking poses and the final result is derived from a weighted consensus score of the two SC2 models. When multiple docking poses are available for a given molecule, the result is based on the pose associated with the highest confidence prediction. The retrieved docking poses are preprocessed by separating the components and applying necessary modifications, including the addition of Gasteiger charges using ADFRSuite (https://ccsb.scripps.edu/adfr/downloads/), to generate the input features for SC2. Evaluation features are extracted from the protein-ligand complexes. As a generalized rescoring method, SC2 reports results without any additional constraints or post-processing steps, such as the conformational optimization of molecules observed in some AI-based docking methods[29, 30].

#### 2.1 Comparing with SC1 on DEKOIS

SC2 was initially benchmarked against its predecessor, SC1, which had previously been evaluated on a subset of the DEKOIS 2.0 dataset, comprising 18 targets as detailed in the original study[4]. For this comparative analysis, the corresponding 18 targets were extracted from our UniDock docking results. These targets, originally a component of DEKOIS 2.0, were isolated from the complete set of 81 targets utilized in subsequent evaluations. Enrichment Factor (EF) values at thresholds of 0.5%, 1%, 2%, and 5% are presented in the radar plot (Figure S4). The plot illustrates the enhanced performance of SC2, with increased coverage indicating an improved capacity for active compound enrichment and smoother curves reflecting greater generalisability across diverse targets. On these 18 targets, SC2 attained EF values of 23.12 (median: 26.57), 19.73 (median: 20.25), 16.04 (median: 16.10), and 8.85 (median: 8.99) at the 0.5%, 1%, 2%, and 5% thresholds, respectively. By comparison, SC1 reported EF values of 15.01 (median: 15.5), 13.78 (median: 15.5), 11.44 (median: 11.16), and 6.89 (median: 5.75) for the same respective thresholds. It is noteworthy that SC1 did not achieve effective virtual screening (VS) results for four targets within the dataset, while SC2 demonstrated effective enrichment for all targets except TS. Furthermore, the second-lowest EF value observed for SC2 was 4.42 at the 0.5% threshold for the CATL target. These marked improvements establish SC2 as a more refined and robust VS method, representing a considerable advancement over SC1.

#### 2.2 SC2 achieved SOTA rescoring performance on classical benchmark

DEKOIS 2.0 and DUD-E are two widely used benchmarks in virtual screening evaluation, distinguished by their active-to-decoy ratios and decoy generation methodologies. DEKOIS 2.0 comprises 81 targets, each with 40 actives and 1,200 structurally diverse decoys, while DUD-E contains 102 targets with a total of 22,886 actives, each paired with approximately 50 computationally generated decoys.

In this section, we present SC2’s performance on both DUD-E and DEKOIS 2.0, comparing it with other published methods (results for these methods are taken from the EquiScore and Surfdock papers [5, 30]). Since DUD-E docking records provide only the top-ranked pose, SC2’s results for DUD-E are presented without considering multiple poses. On DEKOIS 2.0, as shown in Figure 2, SC2 achieved the highest performance, outperforming all other methods at the 0.5%, 1%, and 5% EF levels. Although a direct comparison with all methods is limited by differences in viable docking poses for benchmarking, our analysis—based on released data from EquiScore work [5]—shows that SC2 (EF 0.5% = 21.6) outperformed the recent sota docking method Surfdock (EF 0.5% = 21.0). Additionally, SC2 achieved the best BEDROC score among the compared methods (Fig S1, S2), further highlighting its robust performance. Figure 2 underscores SC2’s top-tier active enrichment, with additional metrics, including AUC-ROC, provided in the Supplementary Material (Fig S1, S2). As SC2 is a consensus model that combines predictions from two independent components(SC2-PB and SC2-PS), the performance of each constituent model is also shown in Figures 2 and 3. On the DUD-E dataset (Figure 3), SC2 again achieved the highest enrichment at all three EF levels, surpassing all other listed methods. These results demonstrate SC2’s superior screening power, its reliability as a hit discovery tool, and its generalisability across diverse targets.

**Fig. 2.**
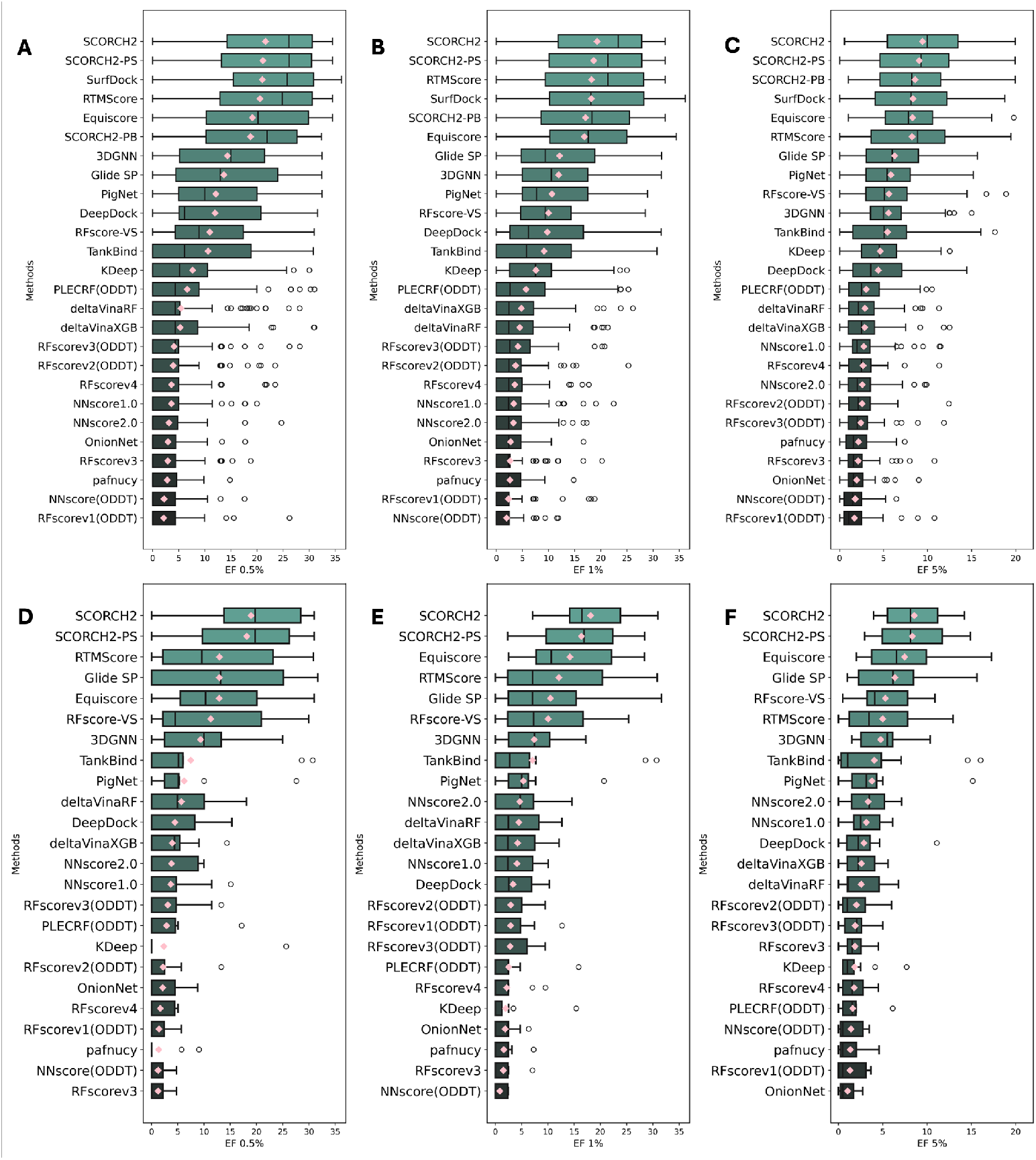
Method evaluation on DEKOIS 2.0 benchmark (data point n=81). Pink diamonds denote the mean EF at a given level; a median line is included in each box-whisker plot. Top row: Model EF on all 81 targets at the 0.5% (A), 1% (B), and 5% (C) levels. Bottom row: Model EF on 11 unseen targets at the 0.5% (D), 1% (E), and 5% (F) levels. SCORCH2-PB: model trained with SC1 and PDBBind data; SCORCH2-PS: model trained with PDBScreen data; SC2: model consensus of SCORCH2-PB and SCORCH2-PS.

**Fig. 3.**
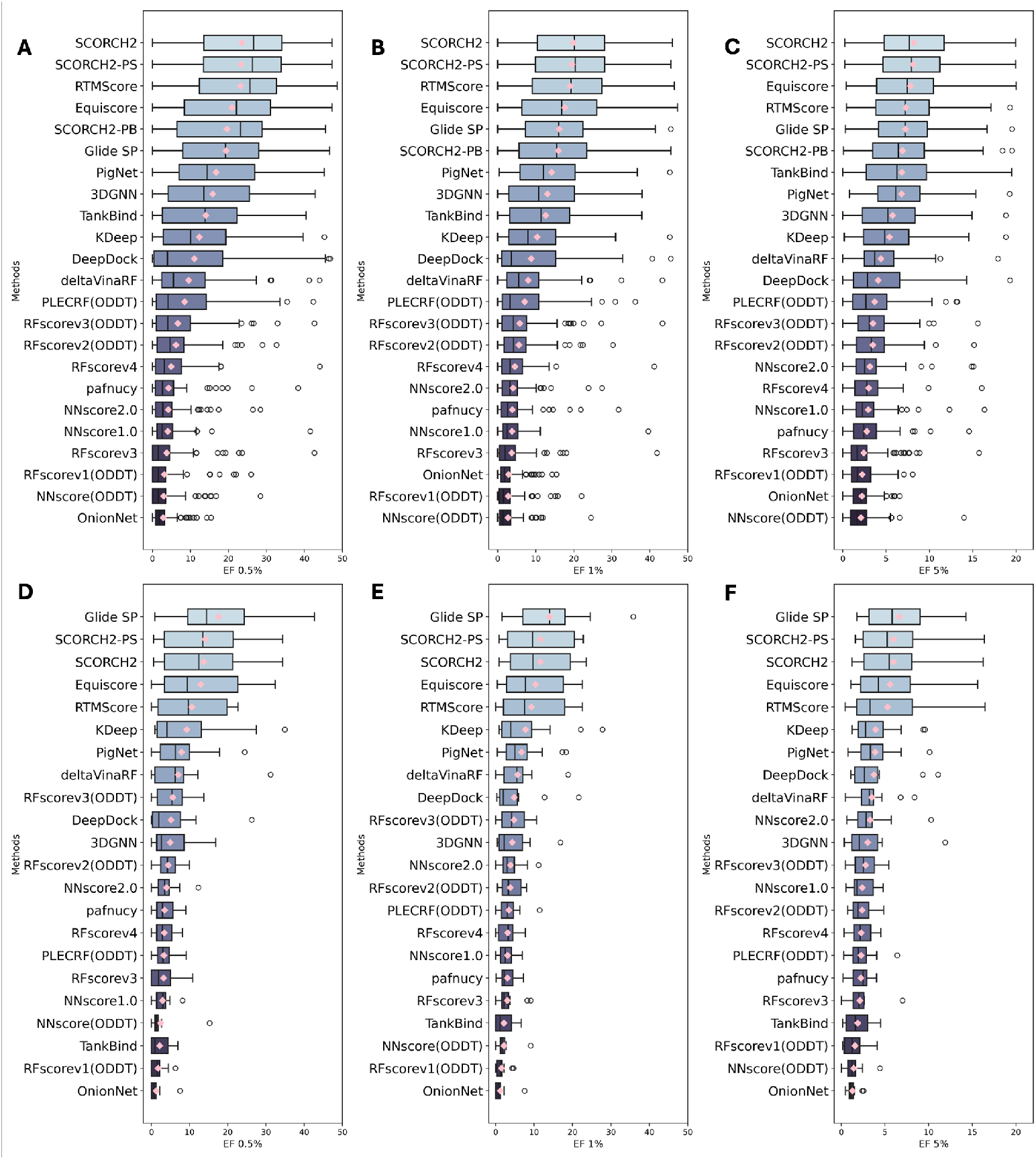
Method evaluation on DUD-E benchmark (data point n=102), the pink diamonds denote mean value for EF at a certain level, where a median line is included in each box-whisker plot. Model EF on all 102 targets at top 0.5% (A), 1% (B), 5% (C) level. Second row, model EF on unseen targets (data point n=12) at top 0.5% (D), 1% (E), 5% (F) level. SC2 result on this plot is without multipose consideration. SCORCH2-PB - model trained with SC1 and PDBBind data, SCORCH2-PS - model trained with PDBScreen data, SC2 - model consensus by SCORCH2-PB and SCORCH2-PS.

#### 2.3 SC2 demonstrates robust transferability to previous unseen targets

Graph Neural Network (GNN)-based methods have exhibited impressive performance [5, 6, 31]; however, their capacity to generalize to previously unseen protein targets remains a critical issue. Prior research [32, 33] has indicated that structural similarity exerts a dominant influence on GNN-based predictions, suggesting that these models may not fully capture the nuances of ligand-receptor interactions. Because GNNs often function without explicit feature engineering, relying instead on purely data-driven learning, their predictions may deviate from established biophysical principles, as ligand binding is fundamentally governed by specific molecular interactions. Consequently, their performance tends to degrade on structurally diverse targets that differ substantially from the training data [5, 16], emphasizing the need for models that more effectively integrate molecular interaction to achieve robust domain adaptation.

In the EquiScore study, the authors identified 11 unseen targets in DEKOIS 2.0 (and 12 in DUD-E) that were absent from the PDBBind database at the UniProt level, a sequence-based classification of protein families. Performance on these targets serves as an indicator of a model’s ability to generalize learned principles to previously unseen targets. To assess the generalisability of SC2, we evaluated its performance on these unseen targets; the results are presented in Figure 2 and Figure 3. It should be noted that SCORCH2-PB incorporates all data from SC1, which includes approximately 2% of data from BindingMOAD[34] and Iridium[35], sources that extend beyond the scope of PDBBind v2020 dataset. A subsequent query of UniProt identifiers from the RCSB PDB revealed six duplicate protein entries within the SCORCH2-PB training data. For the SC2-PS model, our data curation process differed slightly from the original work; the specific steps for data deduplication against these unseen targets are detailed in the Methods section. To maintain a fair comparison, results for SCORCH2-PB are not reported separately for the evaluation of unseen target performance. Tables S2 and S3 provide a detailed listing of the minor data overlap with these unseen targets.

As the result shows, all other methods exhibited substantial performance declines on the 11 unseen DEKOIS targets, whereas SC2 consistently outperformed them, including Glide. On these unseen DEKOIS targets, SC2 achieved an enrichment factor (EF) of 18.94 (median: 19.75) at EF 0.5% and 18.14 (median: 16.53) at EF 1%. On the 12 unseen DUD-E targets, SC2 achieved an EF of 13.71 (median: 12.42) at EF 0.5% and 11.61 (median: 9.76) at EF 1%; in this comparison, Glide exhibited the highest performance, but SC2 still surpassed all other methods. These findings establish that SC2 is highly competitive for screening against unseen protein targets, underscoring its potential as a reliable and innovative tool for virtual screening.

#### 2.4 Case study for SC2 ranking power on Merck FEP benchmark

Ranking represents another critical application for scoring functions (SFs), assessing a model’s capacity to predict compound affinity and its correlation with experimentally determined biological potencies. This capability is particularly important in scenarios such as hit-to-lead optimization, where *in silico* methods like Schrödinger FEP+ [36] are valuable for guiding medicinal chemistry efforts by accounting for minor structural modifications. However, as previously discussed, SC2 is primarily designed as a rescoring method for IBVS, with a focus on leveraging large amounts of unlabeled data. In SC2 development, experimentally determined metrics like binding affinity are not utilized, and all positive data points are treated equivalently. This lack of direct utilization of binding affinity data often leads to reduced performance on ranking or scoring tasks, as previously observed with RTMScore [6]. Subsequent studies, such as GenScore [31], tried to address this limitation by incorporating an adjustable binding affinity term, leading to improved and more balanced performance.

In our investigation, SC2’s features include, among others, energy terms such as electrostatic energy summaries derived from BINANA. This motivated us to investigate the degree to which SC2’s predictions correlate with compound binding affinities. To assess this, we performed a case study using 8 targets from Merck FEP benchmark [37]. The Merck FEP benchmark was created to evaluate the performance of free energy prediction models; it encompasses several pharmaceutically relevant targets and 264 ligands with associated binding affinity data. The ligands (or analogues) for each target share a common core structure but differ in their substituents, simulating realistic drug development scenarios. In our study, we evaluated SC2’s ranking capability on these eight unique target clusters, and Table 1 presents the SC2 results alongside those reported for other methods in the GenScore [31] paper. Evaluations were conducted using the Spearman correlation coefficient (*R*_*s*_) between the SC2 predictions and the experimental Δ*G* values.

**Table 1.**
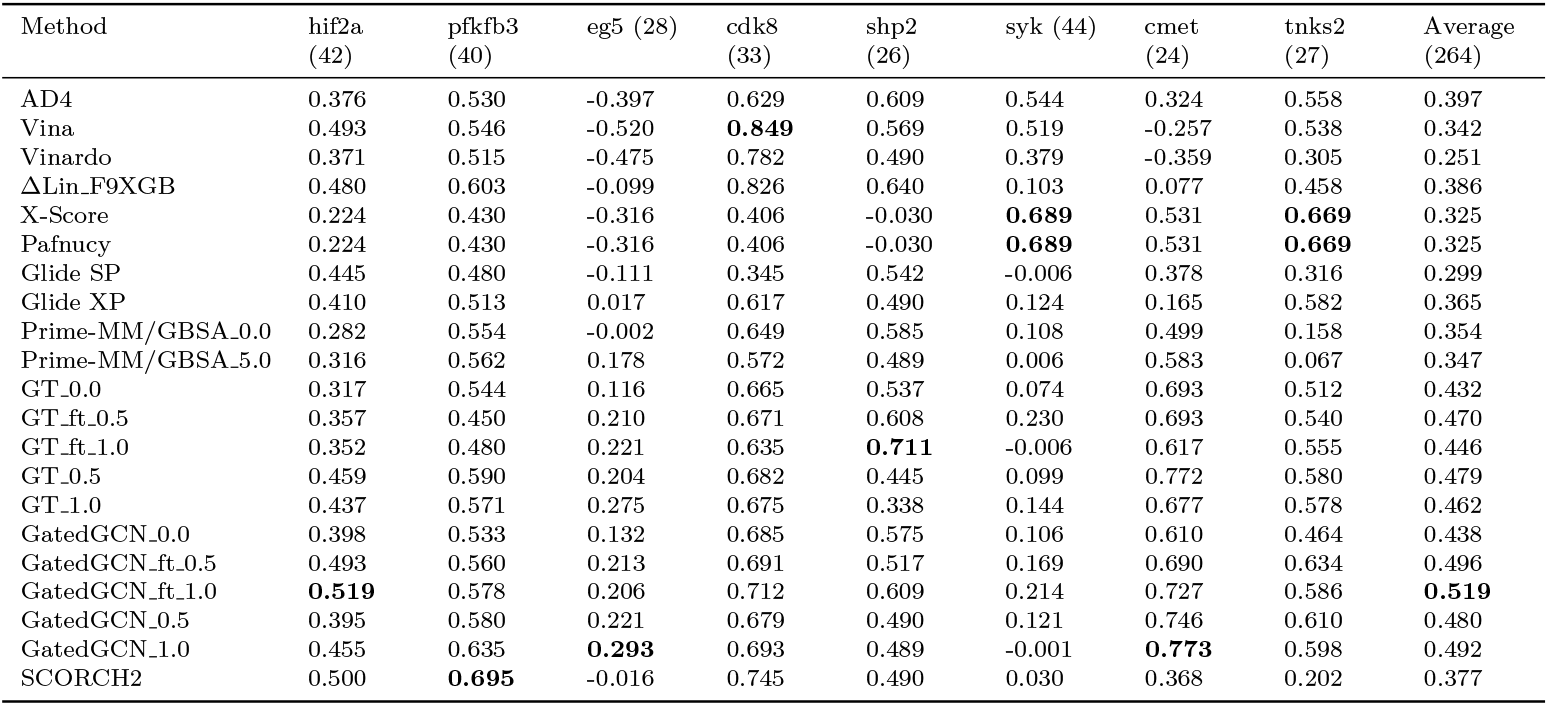
Ranking power comparison between SC2 and other models in terms of Spearman correlation coefficient (*R*_*s*_) on Merck FEP benchmark, eight targets. The best number for each target is bolded.

The results indicate that SC2 exhibits only moderate ranking capability (*R*_*s*_ = 0.377). However, it is crucial to recognize that this performance is attained without incorporating any affinity data during the training process. Despite this inherent limitation, SC2 still outperforms several other methods, including Vina and Glide, in terms of average *R*_*s*_. This observation suggests that SC2 is more sensitive to subtle structural modifications than the aforementioned methods. Nevertheless, due to the ignorance of binding affinity information in the training data, these structural differences are not calibrated against corresponding alchemical energy changes, potentially explaining SC2’s comparatively lower performance in ranking tasks.

#### 2.5 Diverse knowledge patterns can enhance IBVS generalisability

A key limitation in ensemble modelling is that the weakest model in the group can constrain overall performance. Such degradation often stems from feature conflicts or insufficient training [38], and in some cases, ensemble performance may fall short of the best-performing individual model. This issue emerged during early SC2 development.

For instance, SC2-PB underwent extensive optimization, yet consensus performance sometimes declined. SHAP analyses from our RMSD cutoff ablation study (Table S4, Figure S3) suggested that performance loss was not due to poor training but rather local optima tied to dataset scale and distribution. While the two models shared architecture and features, only one consistently outperformed the other—indicating that the weaker model was trapped in a suboptimal region. Though locally optimal, it contributed less effectively to joint predictions, limiting the benefits of consensus.

These findings led to the introduction of the term - Knowledge Pattern (KP), defined as a discrete configuration in parameter space that produces a qualitatively distinct input-output mapping under invariant features. A KP represents more than just parametric variation; it signifies a structural shift in the model’s inductive bias and internal reasoning, which results from exploring different regions of the training space. Therefore, a KP encodes a transformation in how the model interprets and responds to inputs, extending beyond simple output fluctuations.

Our ablation studies revealed that extracting diverse KPs from a single dataset is difficult, even under exhaustive tuning. As shown in Figure S3, differences in top features between models often reflected reweightings rather than distinct combinations, suggesting limited KP diversity in data-isolated settings.

Prior work on consensus docking [39] supports the idea that combining predictions from models with diverse priors improves reliability. To simulate this approach, we incorporated PDBScreen as supplementary training data. PDBScreen, which serves as the training data for Equiscore, is derived from Glide SP docking results, thereby introducing a complementary perspective into the SC2 model. Since different docking protocols generate varied poses due to distinct algorithms and scoring schemes, despite these differences, key interactions—e.g., hydrogen bonding, hydrophobic contacts, and *π*–*π* stacking—tend to be preserved across docking methods. Consequently, binding modes derived from multiple docking methods can act as a valuable enrichment, and help mitigate the limitations of any single docking method while leveraging their collective strengths to enhance predictive performance.

This led to the development of SC2-PS, which was evaluated alongside SC2-PB on DEKOIS 2.0. As shown in Table 2, although SC2-PS outperformed SC2-PB overall, integrating both into a consensus model yielded a performance that surpassed either individual model. This outcome underscores the value of the KP framework in explaining how consensus learning—when rooted in structurally distinct but complementary models—can improve generalization and screening power.

**Table 2.** Performance metrics for various methods. The first eight rows present metrics obtained directly from the respective docking methods themselves. The subsequent rows display the results of rescoring docking poses using other methods: -E indicates EquiScore; -SC2 denotes SCORCH2; and -SC2 PB or -SC2 PS represent rescoring by a single SC2 model (either SCORCH2-PB or SCORCH2-PS, respectively). The best performance for each metric is highlighted in bold.

| Method | AUROC | BEDROC($\alpha = 80.5$) | EF 0.5% | EF 1% | EF 5% |
| --- | --- | --- | --- | --- | --- |
| LEDOCK | 0.6533 | 0.1861 | 6.777 | 5.8557 | 3.6474 |
| SURFLEX | 0.6728 | 0.2202 | 8.3648 | 7.3007 | 3.9998 |
| VINA | 0.6327 | 0.1403 | 5.4641 | 4.5127 | 2.8244 |
| GLIDE | 0.7387 | 0.3855 | 14.6088 | 12.4722 | 6.2995 |
| GOLD | 0.6468 | 0.171 | 5.7502 | 5.3642 | 3.3475 |
| FLARE | 0.6771 | 0.1726 | 6.5404 | 5.5772 | 3.8072 |
| PSOVINA2 | 0.6104 | 0.1295 | 5.3029 | 4.4744 | 2.8269 |
| UNIDOCK | 0.6196 | 0.1326 | 5.3689 | 4.4192 | 2.7546 |
| LEDOCK-E | 0.7986 | 0.3875 | 16.5292 | 14.7716 | 7.1585 |
| SURFLEX-E | 0.8103 | 0.4079 | 15.6913 | 14.4472 | 7.8312 |
| VINA-E | 0.7095 | 0.251 | 10.9728 | 9.3116 | 4.8105 |
| GLIDE-E | **0.821** | 0.460 | 19.095 | 16.832 | 8.274 |
| GOLD-E | 0.7795 | 0.3478 | 15.1629 | 13.0205 | 6.3182 |
| LEDOCK-SC2 | 0.7664 | 0.3890 | 16.7574 | 14.4742 | 7.0611 |
| SURFLEX-SC2 | 0.7677 | 0.4359 | 17.9383 | 15.6992 | 7.6525 |
| VINA-SC2 | 0.7087 | 0.2741 | 13.1910 | 10.1924 | 4.8557 |
| GLIDE-SC2 | 0.748 | **0.549** | **21.622** | **19.264** | **9.405** |
| GOLD-SC2 | 0.7238 | 0.3281 | 15.2142 | 12.3657 | 5.7551 |
| LEDOCK-SC2_PB | 0.7447 | 0.3421 | 14.149 | 12.9241 | 6.5419 |
| SURFLEX-SC2_PB | 0.7606 | 0.4007 | 16.5707 | 14.2806 | 7.092 |
| VINA-SC2_PB | 0.6869 | 0.246 | 11.4885 | 9.2902 | 4.5807 |
| GLIDE-SC2_PB | 0.7384 | 0.4988 | 19.8404 | 17.6569 | 8.7829 |
| GOLD-SC2_PB | 0.7129 | 0.3019 | 13.9679 | 11.1068 | 5.5235 |
| LEDOCK-SC2_PS | 0.7575 | 0.3735 | 16.2736 | 13.7422 | 6.7754 |
| SURFLEX-SC2_PS | 0.7565 | 0.4139 | 16.8971 | 14.87 | 7.2686 |
| VINA-SC2_PS | 0.6973 | 0.2614 | 12.7102 | 9.5705 | 4.6142 |
| GLIDE-SC2_PS | 0.7371 | 0.5287 | 21.0957 | 18.6206 | 9.0385 |
| GOLD-SC2_PS | 0.7104 | 0.3099 | 14.651 | 11.6763 | 5.4249 |
| PSOVina-SC2-max | 0.6737 | 0.4188 | 18.3387 | 15.3603 | 7.0535 |
| PSOVina-SC2-average | 0.6737 | 0.3641 | 14.2506 | 12.6403 | 6.9066 |
| PSOVina-SC2-top1 | 0.7771 | 0.4390 | 18.7251 | 15.8936 | 7.3291 |
| Flare-SC2-max | 0.7202 | 0.4699 | 19.1611 | 16.7133 | 8.4754 |
| Flare-SC2-average | 0.7202 | 0.4211 | 16.9089 | 14.9408 | 7.9089 |
| Flare-SC2-top1 | 0.7771 | 0.4390 | 18.7251 | 15.8936 | 7.3291 |
| Unidock-SC2-max | 0.7119 | 0.4636 | 19.4397 | 17.0351 | 7.8579 |
| Unidock-SC2-average | 0.7119 | 0.4322 | 17.7657 | 15.4353 | 7.7967 |
| Unidock-SC2-top1 | 0.7351 | 0.3813 | 16.7576 | 14.0093 | 6.2451 |
| PSOVina-SC2-unseen-imbalance_weight | 0.7063 | 0.3876 | 16.1696 | 15.2411 | 6.4789 |
| PSOVina-SC2-unseen-balance_weight | 0.7152 | 0.3831 | 16.1699 | 14.1565 | 6.75 |
| Unidock-SC2-unseen-imbalance_weight | 0.7355 | 0.4089 | 15.2948 | 15.1713 | 6.9984 |
| Unidock-SC2-unseen-balance_weight | 0.7443 | 0.4068 | 15.2935 | 14.3028 | 7.0435 |
| Flare-SC2-unseen-imbalance_weight | 0.7350 | 0.4265 | 16.9589 | 15.1779 | 7.5213 |
| Flare-SC2-unseen-balance_weight | 0.7398 | 0.4172 | 16.1553 | 15.8428 | 7.5182 |
| Glide-SC2-unseen-imbalance_weight | 0.7663 | 0.5170 | 18.9423 | 18.1441 | 8.5536 |
| Glide-SC2-unseen-balance_weight | 0.7734 | 0.4977 | 18.0519 | 16.3034 | 8.7150 |

##### 2.5.1 Consensus scheme interpretation

We observed that the SC2 consensus model occasionally underperformed relative to SC2-PS alone on 12 DUD-E targets (Figure 3), likely due to the performance imbalance between SC2-PS and SC2-PB. To evaluate the impact of such imbalance, we conducted two extra experiments on DUD-E and DEKOIS 2.0 datasets. In the first, both models were tested individually on Glide SP docking poses. As shown in Figure S5 (DEKOIS 2.0) and S6 (DUD-E), these stacked bar plots, where all results stem from the same position on the x-axis, illustrate SC2 performance at the EF 0.5%, EF 1% and EF 5% levels. Taking Figure S5 as an example, SC2 consensus results are depicted in forest green with a dark hatch, while SC2-PS and SC2-PB results are shown in dark green and light green, respectively. It is evident that no single model consistently outperforms the others, and results can vary significantly across different targets. Notably, in most cases, the SC2 bar encompasses those of SC2-PS and SC2-PB, suggesting that the consensus model generally achieves a more balanced performance, particularly for top-tier enrichment.

Considering the former experiment is conducted on Glide docking poses and SC2-PS is trained with the data prepared by the same methods, to avoid the energy landscape similarity, the second experiment is done by testing the model separately on DEKOIS 2.0 docking poses provided by another 4 different docking methods in Equiscore paper including LEDOCK[40], SURFLEX[41], VINA[23], and GOLD[42]. The results in Table 2, reveal that in nearly all cases, SC2-PS overall outperforms SC2-PB. However, when considering the entire DEKOIS 2.0 dataset, SC2 generally performs better than SC2-PS when evaluated independently. This suggests that the two models, each with its distinct KP, can complement each other and are less likely to produce conflicting results.

Importantly, SC2 surpassed EquiScore at EF 0.5% across all five docking protocols and on BEDROC for four of them (except GOLD). However, SC2 showed no advantage on AUROC, indicating its strength lies in ranking true actives early rather than improving overall classification. These findings support SC2 as an effective and robust method for successful screening.

##### 2.5.2 Scoring strategy determination

In addition to the poses from EquiScore, we conducted a case study to determine the optimal scoring strategy for SC2. Three scoring patterns were considered:

- Average: the final score is the average of SC2 scores across all available poses for a given molecule.
- Top1: the score corresponding to the top-ranked pose from the docking method is used as the final result.
- Max: the highest SC2 score among all poses is selected, regardless of the original docking rank.

Table 2 presents SC2’s performance on the DEKOIS 2.0 dataset, where additional docking poses were generated using three other methods: PSOVina2[43], Cresset Flare [44–46], and UniDock [47]. Details regarding the docking data preparation can be found in the Methods section. A key finding demonstrated across all eight docking protocols evaluated is that substituting the methods’ native scoring functions with SC2 scores consistently improved virtual screening performance. Informed by the comprehensive results in Table 2, the strategy of utilizing the maximum SC2 score across multiple poses (’max pattern’) was selected for reporting final SC2 results, as it yielded the best outcomes. These findings further underscore SC2’s broad applicability as a generalised rescoring framework.

##### 2.5.3 Consensus weight determination

After confirming the optimal scoring strategy, we further investigated the determination of consensus weights for the final output. Previous results demonstrated that model consensus improves generalisability across diverse targets in the DEKOIS 2.0 dataset. However, the optimal contribution of each model to the final consensus prediction remains an open question. Here, consensus weight refers not to trainable parameters but to the proportion of each model’s contribution to the final score.

To assess the impact of different weighting strategies, we evaluated SC2 on 11 unseen targets from the DEKOIS 2.0 dataset, using docking poses generated by Glide SP and three additional docking protocols. Two strategies were examined:

- Balanced weight, where the final score was the average of SC2-PS and SC2-PB predictions, remains identical in SC1.
- Imbalanced weight, where SC2-PS contributed 70% and SC2-PB 30% to the final score, the percentage is empirically determined.

As shown in Table 2, the balanced strategy yielded improved VS performance at the EF 5% level, while the imbalanced strategy achieved better enrichment at EF 0.5% and EF 1% level. Since early enrichment is more critical in practical compound selection, the imbalanced approach offers better applicability. Furthermore, maximizing generalisability may come at the cost of reduced per-target performance. We therefore recommend the imbalanced consensus weighting to better balance sensitivity and generalisability in real-world virtual screening applications.

#### 2.6 Benchmarking SC2 on new VSDS-vd dataset

VSDS-vd is a recently introduced benchmark for systematic evaluation of AI-powered and physics-based docking methods [48]. It includes three subsets—*TrueDecoy, RandomDecoy*, and *MassiveDecoy*—designed to assess docking accuracy and virtual screening (VS) performance under increasing complexity. *TrueDecoy* comprises 147 targets with experimentally validated actives and inactives. *RandomDecoy* introduces decoys sampled from commercial libraries, while *MassiveDecoy* extends to 7 million decoys for large-scale VS. All subsets were curated from BindingDB [49], ChEMBL [50], PubChem [51], and PDB, ensuring target diversity with an active-inactive ratio of 1:40.

Their study evaluates four AI-based docking methods (CarsiDock, KarmaDock, DiffDock, FlexPose), four physics-based tools (Glide, LeDock, rDock [52], Surflex), and two rescoring functions (RTMScore, EquiScored). In our study, we also evaluated SC2 on the VSDS-vd dataset. However, due to the commercial nature of Glide and the absence of docked poses, SC2’s performance on Glide docking poses cannot be further assessed. In that case, SC2 was evaluated using Flare docking poses, which previously showed superior performance. Docking and processing details are described in the Methods section. To manage computational cost, we focus on the two most challenging subsets: *TrueDecoy* and *TrueDecoy gap*.

Results are shown in Table 3. In general, rescoring with SC2 or RTMScore improves virtual screening outcomes, especially in early enrichment. It is important to note that Flare provides multiple scoring metrics; for this evaluation, only the LF VSscore was used. As a newly established benchmark, VSDS-vd introduces more challenging scenarios and a broader range of targets, making it a valuable resource for evaluating virtual screening methods. Future studies could further explore the dataset’s distribution and potential hidden biases. In conclusion, our findings demonstrate that SC2 is a generalized and effective rescoring approach, significantly improving active compound enrichment in virtual screening.

**Table 3.**
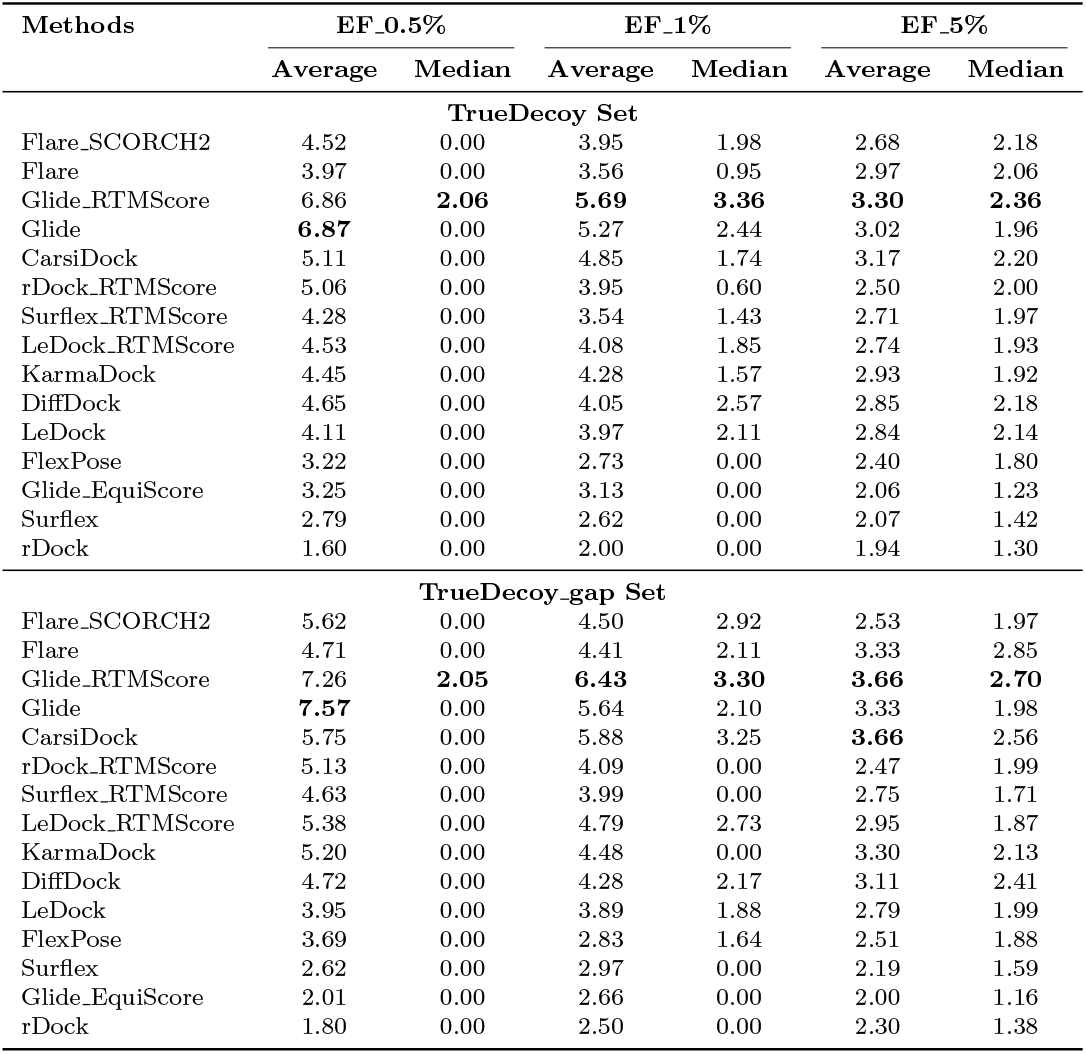
Enrichment Factors (EF) for Various Docking Methods on the TrueDecoy and TrueDecoy gap Sets. The highest number is bolded.

| Methods | EF_0.5% |  | EF_1% |  | EF_5% |  |
| --- | --- | --- | --- | --- | --- | --- |
|  | Average | Median | Average | Median | Average | Median |
| **TrueDecoy Set** |  |  |  |  |  |  |
| Flare_SCORCH2 | 4.52 | 0.00 | 3.95 | 1.98 | 2.68 | 2.18 |
| Flare | 3.97 | 0.00 | 3.56 | 0.95 | 2.97 | 2.06 |
| Glide_RTMScore | 6.86 | **2.06** | **5.69** | **3.36** | **3.30** | **2.36** |
| Glide | **6.87** | 0.00 | 5.27 | 2.44 | 3.02 | 1.96 |
| CarsiDock | 5.11 | 0.00 | 4.85 | 1.74 | 3.17 | 2.20 |
| rDock_RTMScore | 5.06 | 0.00 | 3.95 | 0.60 | 2.50 | 2.00 |
| Surflex_RTMScore | 4.28 | 0.00 | 3.54 | 1.43 | 2.71 | 1.97 |
| LeDock_RTMScore | 4.53 | 0.00 | 4.08 | 1.85 | 2.74 | 1.93 |
| KarmaDock | 4.45 | 0.00 | 4.28 | 1.57 | 2.93 | 1.92 |
| DiffDock | 4.65 | 0.00 | 4.05 | 2.57 | 2.85 | 2.18 |
| LeDock | 4.11 | 0.00 | 3.97 | 2.11 | 2.84 | 2.14 |
| FlexPose | 3.22 | 0.00 | 2.73 | 0.00 | 2.40 | 1.80 |
| Glide_EquiScore | 3.25 | 0.00 | 3.13 | 0.00 | 2.06 | 1.23 |
| Surflex | 2.79 | 0.00 | 2.62 | 0.00 | 2.07 | 1.42 |
| rDock | 1.60 | 0.00 | 2.00 | 0.00 | 1.94 | 1.30 |
| **TrueDecoy-gap Set** |  |  |  |  |  |  |
| Flare_SCORCH2 | 5.62 | 0.00 | 4.50 | 2.92 | 2.53 | 1.97 |
| Flare | 4.71 | 0.00 | 4.41 | 2.11 | 3.33 | 2.85 |
| Glide_RTMScore | 7.26 | **2.05** | **6.43** | **3.30** | **3.66** | **2.70** |
| Glide | **7.57** | 0.00 | 5.64 | 2.10 | 3.33 | 1.98 |
| CarsiDock | 5.75 | 0.00 | 5.88 | 3.25 | **3.66** | 2.56 |
| rDock_RTMScore | 5.13 | 0.00 | 4.09 | 0.00 | 2.47 | 1.99 |
| Surflex_RTMScore | 4.63 | 0.00 | 3.99 | 0.00 | 2.75 | 1.71 |
| LeDock_RTMScore | 5.38 | 0.00 | 4.79 | 2.73 | 2.95 | 1.87 |
| KarmaDock | 5.20 | 0.00 | 4.48 | 0.00 | 3.30 | 2.13 |
| DiffDock | 4.72 | 0.00 | 4.28 | 2.17 | 3.11 | 2.41 |
| LeDock | 3.95 | 0.00 | 3.89 | 1.88 | 2.79 | 1.99 |
| FlexPose | 3.69 | 0.00 | 2.83 | 1.64 | 2.51 | 1.88 |
| Surflex | 2.62 | 0.00 | 2.97 | 0.00 | 2.19 | 1.59 |
| Glide_EquiScore | 2.01 | 0.00 | 2.66 | 0.00 | 2.00 | 1.16 |
| rDock | 1.80 | 0.00 | 2.50 | 0.00 | 2.30 | 1.38 |

**Table 4.** Performance metrics of various scoring functions across DEKOIS 2.0 and DUD-E hard targets at EF 0.5% and EF 1%. NA denotes the data not mentioned in the former publication. The highest number is bolded.

| Target | Glide SP | EquiScore | RTMScore | Karmadock_raw | SCORCH2 |
| --- | --- | --- | --- | --- | --- |
| EF 0.5% |  |  |  |  |  |
| HIV1RT | 18.30 | 16.43 | 14.09 | 0.00 | 9.39 |
| HSP90 | 4.34 | 5.05 | 25.99 | 20.22 | 25.99 |
| QPCT | 0.00 | 5.17 | 31.00 | 25.79 | 4.43 |
| SARS-HCoV | 4.40 | 0.00 | 0.00 | 6.15 | 0.00 |
| SIRT2 | 4.36 | 5.08 | 4.35 | 0.00 | 4.35 |
| Average | 6.28 | 6.35 | **15.09** | 10.43 | 8.83 |
| MCR | 25.12 | 16.23 | 32.46 | NA | 32.65 |
| LKHA4 | 18.83 | 26.01 | 30.21 | NA | 31.11 |
| MMP13 | 25.20 | 37.71 | 28.87 | NA | 36.92 |
| KITH | 22.60 | 22.60 | 22.60 | NA | 19.59 |
| FNTA | 2.43 | 13.63 | 1.10 | NA | 9.82 |
| Average | 18.84 | 23.24 | 23.05 | NA | **26.02** |
| EF 1% |  |  |  |  |  |
| HIV1RT | 9.85 | 10.96 | 7.58 | 0.00 | 7.58 |
| HSP90 | 2.33 | 10.11 | 23.33 | 20.22 | 23.33 |
| QPCT | 0.00 | 5.17 | 31.00 | 25.79 | 4.77 |
| SARS-HCoV | 2.37 | 0.00 | 0.00 | 5.59 | 2.37 |
| SIRT2 | 2.35 | 7.62 | 2.34 | 0.00 | 2.34 |
| Average | 3.38 | 6.77 | **12.85** | 10.32 | 8.078 |
| MCR | 19.53 | 14.88 | 20.29 | NA | 26.05 |
| LKHA4 | 16.99 | 22.24 | 26.43 | NA | 24.45 |
| MMP13 | 20.71 | 37.41 | 20.37 | NA | 31.65 |
| KITH | 22.60 | 22.60 | 22.60 | NA | 20.34 |
| FNTA | 2.06 | 11.79 | 1.40 | NA | 9.16 |
| Average | 16.37 | 21.78 | 18.61 | NA | **22.33** |

#### 2.7 SC2 provides interpretable predictions aligned with chemical principles

One of the key limitations in applying machine learning to drug discovery is the lack of interpretability, often referred to as the black-box problem. This issue obscures how models make decisions and which features drive their predictions. While SC2 does not include explicit structural information, which makes atom-level attention and similar interpretations not applicable, it enhances global interpretability by offering clear explanations that align with established chemical principles.

We then quantify feature importance using SHAP, which estimates each feature’s marginal contribution to a prediction [27]. Figure 4 shows SHAP beeswarm plots for SC2-PB and SC2-PS models, revealing that molecular contact descriptors and energy terms dominate the predictions, reflecting chemical reasoning. For example, in SC2-PS, top features include `2.5(HD,OA)`, representing hydrogen bonds between polar hydrogens and oxygen acceptors, and `4.0(A,C)`, indicating hydrophobic contacts involving aromatic and aliphatic carbons. Higher values of `2.5(HD,OA)` and more negative `ElSum(HD,OA)`—reflecting favourable hydrogen bonding—correlate with stronger predicted binding, consistent with chemical principles. In contrast, `2.5(C,HD)`, which captures interactions between carbon atoms and hydrogen donors, shows a “more-to-worse” trend, aligning with the low electronegativity of carbon and its poor hydrogen bond acceptor capability. These results support the validity of SC2’s feature design and demonstrate that its predictions are grounded in chemical reasoning.

**Fig. 4.**
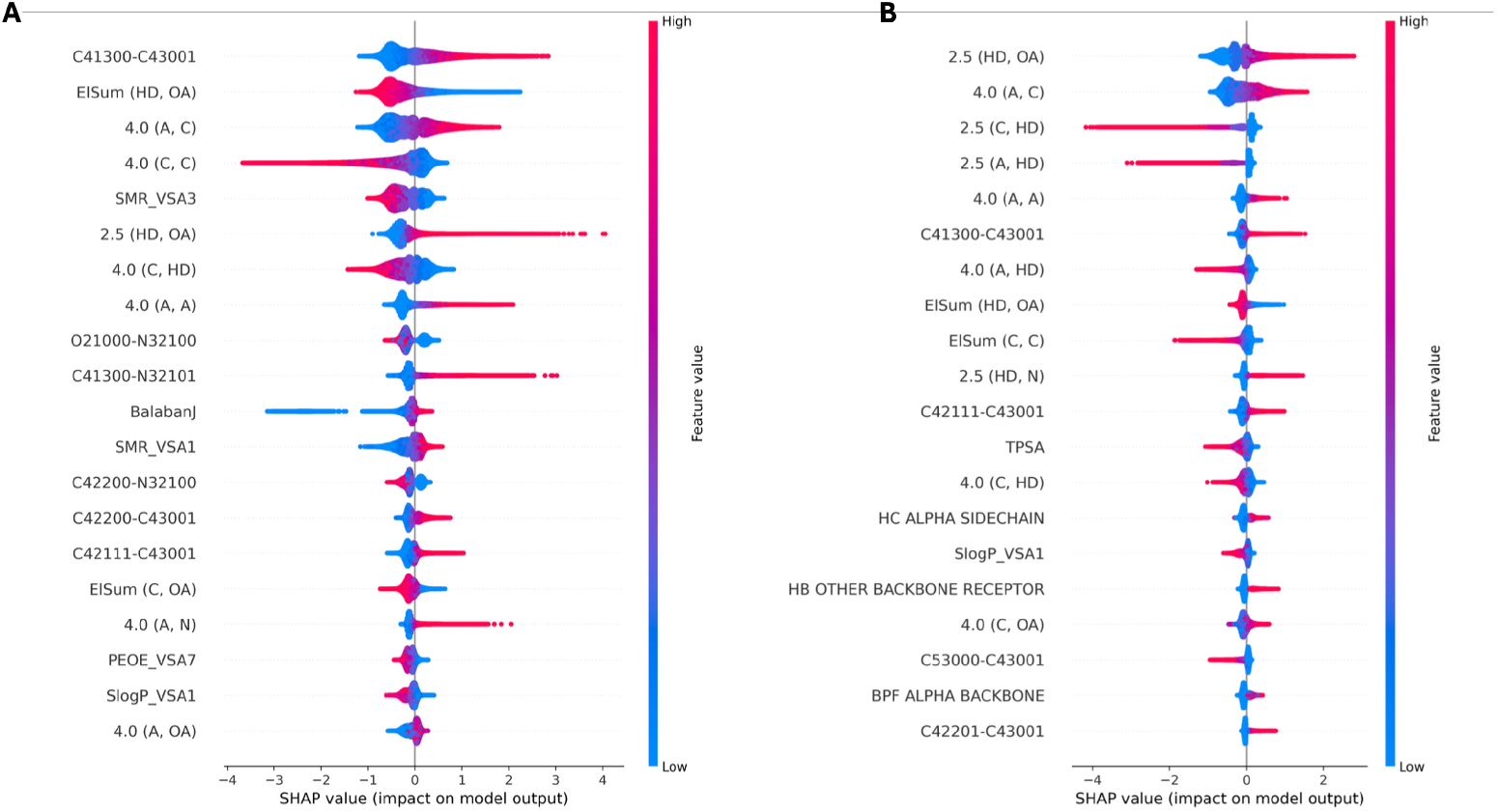
SHAP value beeswarm plot visualization. The color of the figures indicates the value of each feature (blue for lower numbers and red for higher numbers) while the thickness of the line corresponds to the number of samples. Features are ranked by importance with the most important feature at the top. A, SHAP feature ranking of SC2-PB; B, SHAP feature ranking of SC2-PS.

Beyond global feature importance, SC2 also enables ligand-level interpretation of binding modes. Because its features explicitly incorporate specific molecular interactions, SC2 can offer insight into whether a particular interaction promotes or impairs binding, offering valuable insights for drug discovery.

To illustrate this capability, we analyzed two Spleen Tyrosine Kinase (SYK, target PDB ID: 4PV0) inhibitors from the Merck FEP benchmark: CHEMBL3264995 (4-[(3-8-[(3,4-dimethoxyphenyl)amino]imidazo[1,2-a]pyrazin-6-ylbenzoyl)amino]benzoic acid) and CHEMBL3265032 (ENTOSPLETINIB), with experimental ΔG of − 12.38 and − 11.06 kcal/mol, respectively. Although both compounds share a core imidazo[1,2-a]pyrazine scaffold, they differ in distal interactions: CHEMBL3264995 forms hydrogen bonds with Asn457 and a water molecule, while CHEMBL3265032 compensates with a rotated indazole that forms a new bond with Asp512. Notice that our analysis is conducted based on the data in FEP benchmark so the binding detail can be different from the reported work [53].

Figure 5 displays 2D interaction plots and SHAP waterfall plots, which reveal the influence of distinct interaction profiles on SC2 predictions. For CHEMBL3265032, the absence of terminal benzoic acid precludes hydrogen bonding with Asn457, resulting in negative SHAP contributions from feature `2.5(HD,OA)`. In contrast, CHEMBL3264995 exhibits stronger predicted binding potential, partly attributed to additional hydrophobic contacts, as reflected by higher values of the feature `4.0(A,C)` and its positive SHAP contribution. These observations underscore SC2’s ability to infer binding mode differences from meaningful interaction features, offering chemically coherent and interpretable outputs.

**Fig. 5.**
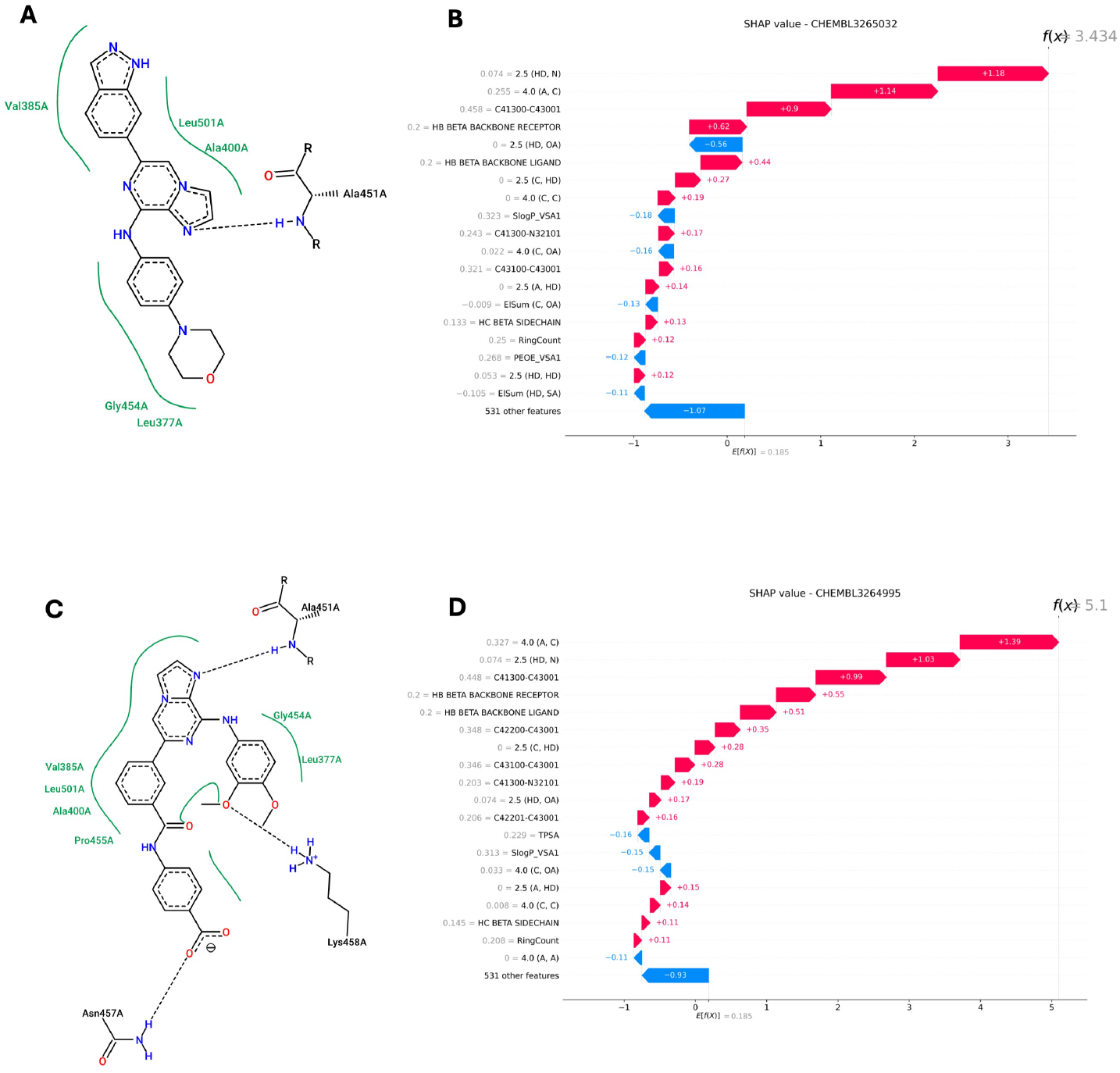
Analysis of molecular interactions and feature importance for two SYK inhibitors using 2D interaction diagrams and SHAP waterfall plots. (A; B) Analysis for CHEMBL3265032: 2D interaction diagram (A) and SHAP waterfall plot (B). (C; D) Analysis for CHEMBL3264995: 2D interaction diagram (C) and SHAP waterfall plot (D). In the SHAP waterfall plots, the color of each bar indicates the value of the corresponding feature (blue for lower values and red for higher values). Features are ranked by their contribution to the model’s prediction, with the most important feature at the top.

#### 2.8 Is physics all you need for virtual screening?

The concern regarding the physical plausibility of deep learning models has garnered significant attention in recent years [48]. Appropriate exposure to physical features may lead to improved pose sensitivity and unlock a model’s potential for novel targets [54]. In this study, we demonstrate that physics or interaction-inspired models can achieve robust virtual screening performance, even on previously unseen targets. However, this also prompts us to consider an alternative question: Is physics all you need for successful virtual screening?

The challenge of screening against unseen targets in IBVS is critical. However, it is equally important to acknowledge that IBVS also continues to face inherent limitations stemming from the characteristics of the targets themselves. We refer to a separate group of targets from previous benchmarks that exhibit these limitations as “hard targets”, defined as targets for which prior research has failed to yield satisfactory screening results due to factors such as receptor flexibility, missing or altered binding subsites, predominantly hydrophobic binding pockets, limited ligand diversity, or high decoy similarity to actives. We reviewed the retrospective screening experiments reported in the DUD-E and DEKOIS 2.0 datasets and identified several such challenging targets. The hard targets from DEKOIS 2.0 include HIV1RT, HSP90, QPCT, SARS-CoV, and SIRT2, which were reported in the initial work. For the DUD-E dataset, we selected an equal number of representative targets with the worst screening performance within each target class, including MCR, LCK, MMP13, KITH, and FNTA. All SC2 results were obtained from prior evaluations, and the data presented in the accompanying chart correspond to the EF 0.5% and 1% levels.

Our results indicate that although SC2 relies solely on general molecular descriptors and omits explicit structural information, it nonetheless achieves competitive performance. However, when dealing with targets that pose structural challenges, reliance on biophysical features introduces inherent limitations. For instance, in cases where traditional docking methods underperform due to high decoy similarity—such as MMP13 in DUD-E—SC2 and EquiScore outperform Glide, highlighting their greater capacity for ligand enrichment under such conditions. Conversely, for targets characterized by shallow, hydrophobic, or highly flexible binding pockets, all evaluated methods showed varying degrees of performance decline compared to their overall dataset averages. This highlights the implicit yet significant impact of target-intrinsic properties on screening outcomes. We therefore emphasize the importance of recognizing these potential challenges during prior evaluation to ensure reliable and optimal virtual screening performance.

#### 2.9 When native scoring is “good enough”: could alternative SFs still improve results?

As we discussed in the last section, alternative scoring functions (SFs) can be influenced by target-intrinsic problems. Intuitively, we also investigated whether alternative SFs could outperform the native one when it already yields a “good enough” outcome. We used the Glide score as an example, examining marginal performance by setting challenging thresholds. Specifically, for both DUD-E and DEKOIS, we set enrichment numbers of 20 for EF 0.5%, 15 for EF 1%, and 8 for EF 5%. Figures S7 and S8 present results only for targets where the native Glide score met these hard thresholds. Notably, SC2 outperformed the native Glide score in more than half of the cases. This reinforces the value of rescoring docked poses, even when the initial scoring produces ideal results.

## 3 Discussion

In conclusion, SC2 builds upon SC1 by adopting a more comprehensive and diverse approach to enhance IBVS performance and generalisability, leading to a more reliable, robust, and interpretable framework. By focusing on generalizable interaction patterns, SC2 demonstrates strong applicability to novel targets and highlights the value of molecular interaction-driven approaches in VS scenarios. This suggests that physics-driven methods with well-structured domain knowledge remain competitive with AI-driven and data-driven methods. Additionally, SC2 explores the integration of different KP to improve performance across targets within identical feature constraints, further underscoring the potential of consensus modelling.

Nevertheless, we acknowledge that physics-based approaches, while interpretable and grounded in biophysical principles, are more susceptible to performance degradation in structurally challenging targets, primarily due to their reliance on accurate docking poses. This highlights the importance of further investigating the dependency of IBVS models on docking quality. On the other hand, recent AI-based virtual screening models [7, 55], particularly those leveraging large-scale pretrained encoders, may demonstrate promising robustness against challenging targets, potentially due to their independence from molecular docking and the ability to avoid the associated pose prediction errors. However, whether such models can reliably capture drug-like molecular characteristics—beyond statistical or pattern-based representations—remains an open question.

We believe that the future of virtual screening may lie in the thoughtful integration of AI-based and physics-based methodologies, leveraging their complementary strengths: the scalability of AI with the interpretability and domain fidelity of physical modelling. We hope this work inspires further exploration of hybrid approaches and contributes to the development of more robust, generalizable, and mechanistically grounded tools for accelerating drug discovery.

### 4 Methods

#### 4.1 Data

SC2 builds upon its predecessor SC1 by incorporating all previous data derived from PDBBind[56], BindingMOAD[34], and Iridium[35], along with the remaining entries from the PDBBind v2020 dataset included for SC2-PB training. Data from SC1, with proteins processed uniformly using MGLTools, remained unaltered while preserving relevant cofactors such as metal ions. The original decoys from SC1 were discarded, and a new decoy set was generated via Tocodecoy[25] for SC2. For the additional SC2-PB data, all HET groups were excluded and receptors were treated as rigid. Data splitting followed the bias control strategy of LP-PDBBind [57], which aims to improve generalization by minimizing sequence similarity between training and other data groups. SC2 adopted this strategy with the modification described below.

For SC2-PS training, the PDBScreen dataset was employed. PDBScreen compiles protein-ligand complexes from the Protein Data Bank (PDB), retaining only those with resolution better than 2.5 Å and excluding complexes containing endogenous molecules, crystallographic additives, or covalent binders. The dataset includes redocked poses to increase the representation of near-native poses (RMSD ≤ 2 Å) and low-energy alternatives. Negative samples are generated through cross-docking, in which ligands are docked against ten randomly selected non-interacting proteins. To enhance the complexity of negative samples and mitigate artificial enrichment bias, DeepCoy[26] generates decoys that maintain similar physicochemical properties but exhibit distinct chemical structures, and the top five decoys are selected based on shape similarity to the crystal ligand. The authors of PDBScreen performed deduplication by removing training entries associated with UniProt IDs overlapping with those in external test sets (DUD-E and DEKOIS 2.0), and reallocating them to an internal test set.

##### Data Split Adjustments

###### 1. External Validation

SC2 eliminates the internal validation set, allocating more data to training. The LP-PDBBind training and testing sets were combined with all SC1 data to form a new, comprehensive training set. The LP-PDBBind validation set was retained as the new test set, providing a smaller, dissimilar subset to monitor overfitting. SC2 is validated solely on common benchmarks.

###### 2. PDBScreen Splitting

We adopted the original data split of PDBScreen from the EquiScore study[5], which partitions the dataset into low-quality (LQ) and high-quality (HQ) subsets, both of which were included in SC2. Docking poses and corresponding binding pockets were extracted from binary-stored files. To avoid data leakage, we merged the original training and test sets for model development while retaining the original internal validation set as the new test set for model selection and performance monitoring. All ligand and pocket structures were processed using ADFRSuite to assign charges. To ensure no target overlap with external benchmarks, we further removed any proteins in the test set sharing UniProt IDs with those in DUD-E or DEKOIS2.0, and Table S1 listed all the removed entries in the PDBScreen dataset.

##### Data Curation

###### 1. SC2-PB Training Data Preparation with UniDock

UniDock[47], a GPU-accelerated variant of AutoDock Vina, was employed to redock crystallized ligands. The detailed mode was applied for ligands, while the fast mode was used for decoys. Molecules were converted from `mol2` to `PDBQT` format using ADFRsuite, with Gasteiger charges added via `prepare_ligand`. Receptors were processed using `prepare_receptor`; those failing conversion were preprocessed with pdb2pqr before a second attempt. Successfully converted receptors were included in the training data, while persistent failures were excluded. The binding center was defined as the native pose’s center of mass, with a docking box of 30 Å *×* 30 Å *×* 30 Å. Crystallized poses were also retained as positive samples.

###### 2. Successful Docking Pose Definition

SC2 set three different thresholds for active tolerance. Empirically, poses with RMSD ≤ 2 Å are considered successful for reproducing native poses. Additional cutoffs of 2.5 Å and 3 Å denote moderate and maximized tolerance, respectively. RMSD computation was performed using the Unidock toolkit, ignoring all hydrogens (polar or non-polar). Filtered poses were retained as positive training data.

###### 3. Decoy Generation with Tocodecoy

Tocodecoy, an improved variant of DeepCoy, was used to generate diverse decoys simulating realistic VS conditions. Up to 150 decoys per PDB ID were randomly selected and docked using the same protocol as for ligands. Decoys were not generated for ligands exceeding 700 Dalton or containing more than 32 rotatable bonds.

###### 4. Negative Training Samples

Training with dissimilar decoys mimics real-world scenarios in virtual screening tasks [58]. However, it introduces the potential risk of manual bias, as these decoys are intentionally designed to exhibit significant structural differences from true binders. This structural disparity often leads to variations in properties such as charge distribution, which the model may exploit, potentially inflating test set performance. To mitigate this, we followed recommendations from previous studies [54] and incorporated redocked crystal ligand poses with RMSD ≥ 4 Å as additional negative training samples. This approach helps establish clearer decision boundaries and reduces ambiguity in model predictions.

###### 5. Additional Docking Protocols

For DEKOIS 2.0, official 3D conformations were used with search space confined to the binding pocket. The docking box was defined as an 8 Å buffer around the crystallized ligand’s center of mass (computed via Pybel). PSOVina2 and UniDock shared identical settings, with PSOVina2 exhaustiveness set to 32 and detailed mode for Unidock. Flare was run in reference mode with automatically inferred binding boxes. All methods retained up to 10 poses for SC2 rescoring.

### 4.2 Model

#### SC2 introduced more molecular descriptors

SC1 incorporated 492 distinct features derived from three established methods: `BINANA`[59], `ECIF`[60], and `Kier-Flexibility`[61]. An illustrative example of a molecular interaction is presented in Figure 1 (`PDBID: 1AFK`, `HET Identifier: PAP`, `Name: 3’-PHOSPHATE-ADENOSINE-5’-DIPHOSPHATE`), visualized using the `BINANA` web server (https://durrantlab.pitt.edu/binana/).

`BINANA` identifies ligand-receptor interactions based on distance cutoffs of 2.5 Å or 4 Å, capturing a range of binding features such as hydrogen bonds, *π*–*π* stacking, and electrostatic interactions. The electrostatic interaction energy is calculated using Equation 1, where *V*_(*a,b*)_ represents the summed electrostatic interaction energy between atom types *a* and *b*. Here, 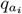 and 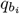 denote the partial atomic charges of atoms of types *a* and *b*, respectively, and 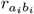 is the interatomic distance. Partial charges are assigned according to AutoDock atom types.

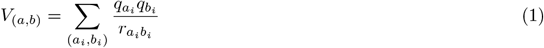

In parallel, `ECIF` performs atom-level interaction counting within defined distance thresholds, retaining finer-grained structural information. This complements `BINANA`, which captures broader interaction patterns but lacks atom-level resolution. In SC2, all features and extraction methods from SC1 were preserved for consistency and further enhancement.

To comprehensively represent the ligand–receptor complex, features can be broadly categorized into three components: ligand, receptor, and interaction. Since the interaction component has already been thoroughly modelled, receptor-level descriptors were excluded from SC2 for two primary reasons:

1. Previous studies have shown that protein features contribute minimally to model performance[62].
2. In virtual screening scenarios, decoys and active ligands are associated with the same receptor, rendering protein descriptors redundant.

Although receptor features were excluded, ligand-specific information remained underutilized in SC1. Prior research has demonstrated that incorporating ligand properties can enhance structure-based predictive models[63]. Moreover, the ligand itself may be regarded as a proxy for the binding mode. Based on this rationale, SC2 introduced 59 additional ligand descriptors computed using `rdkit.ML.Descriptors`. These features primarily capture 1D and 2D molecular characteristics, including topological indices (e.g., `FractionCSP3`, `ChiXv`, `BalabanJ`, `BertzCT`, `HallKierAlpha`) and structural properties (e.g., `HeavyAtomMolWt`, `RingCount`).

Notably, although interaction fingerprints such as `ECIF` were employed, SC2 maintained a design principle of information neutrality. That is, interaction features are not directly mapped to discrete ligand substructures and can arise anywhere within the molecule. This approach mitigates the risk of substructure-specific bias and ensures that model decisions are not confounded by localized ligand motifs. Consequently, SC2 preserves its objective of unbiased and generalizable predictions across diverse ligand chemotypes.

#### 4.3 Why XGBoost?

SC1 also adopted an ensemble strategy, with final predictions obtained by averaging outputs from multiple base learners. Specifically, SC1 combined a single XGBoost model with 15 instances each of two types of neural networks—wide-and-deep and feedforward architectures—to form a consensus ensemble. SC2 retains this ensemble philosophy but simplifies the architecture by employing only two distinct XGBoost models for prediction.

This modification is motivated by empirical evidence. First, among all individual models in SC1, the XGBoost model achieved the highest evaluation performance. Second, a recent systematic benchmark study[64] comparing a wide range of machine learning and deep learning models on tabular data concluded that tree-based methods remain superior in such settings. Notably, XGBoost demonstrated the best overall performance after extensive hyperparameter optimization.

An additional motivation relates to the nature of the curated features in SC2, particularly the ECIF interaction descriptors. As molecular structures are inherently flexible, many ECIF features tend to be sparsely populated, with a substantial proportion being uninformative (i.e., zero-valued). The same study[64] further evaluated model robustness under varying proportions of uninformative features and found that tree-based models, especially XGBoost, maintained superior predictive stability and performance.

#### 4.4 Training details

In SC2, `XGBoost` serves as the sole model architecture. While most machine learning models benefit from data normalization for improved training efficiency, tree-based models like `XGBoost` are naturally resilient to unnormalized data and its skewness. Therefore, SC2 employs `MaxAbsScaler()` from `scikit-learn` to normalize the features, preserving their original distribution and inherent skewness. This approach is based on the hypothesis that retaining the true distribution of features allows the model to better capture the underlying patterns associated with positive samples.

Parameter tuning is crucial for optimizing the performance of tree-based models. Compared to the `gp minimize` method used in SC1, the `Optuna` library [65] provides a more flexible and efficient framework for hyperparameter optimization. In SC2, Optuna integrates a comprehensive set of hyperparameters into an objective function for systematic exploration; the encapsulated parameters for SC2 training are listed in Table S5. For SC2-PS and SC2-PB, each `XGBoost` model undergoes 100 epochs of parameter searching with a maximum of 2K steps per iteration to preview the performance in the current setting. An early stopping criterion of 50 steps is employed to prevent overfitting. Notably, SC2 directly trains on the imbalanced dataset without additional manual data augmentation of the prepared training set or PDBScreen data. In imbalanced datasets, where the majority class dominates, models often exhibit a bias toward this class. To address this, loss compensation is implemented by increasing the weight of positive samples, as described in equation 2, where *y*_train_ denotes the label of each training sample.

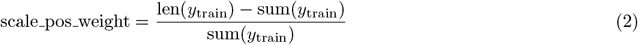

Balancing the weight of actives alone does not guarantee optimal performance. A recent study emphasizes the need to prioritize metrics that focus on correctly classifying actives in virtual screening [66]. In this work, we target the maximization of AUCPR (Area Under the Precision-Recall Curve) for optimization. AUCPR is particularly suited for imbalanced datasets, as it evaluates the model’s ability to identify positive instances, which are typically the minority class.

Once the optimal parameters are identified, the final model undergoes continuous training with a cap of 50K steps. The training process persists until the maximum step is reached or overfitting is detected, as signalled by the early stopping mechanism triggered after 100 rounds. Loss compensation is handled by amplifying the weight for proportional adjustment and the loss function is set as weighted binary cross-entropy as equation 3, where *y*_*i*_ is the true label (0 or 1) for the *i*-th sample, *p*_*i*_ is the predicted probability for class 1 for the *i*-th sample, and *N* is the total number of samples and *w*_*i*_ is the multiplicative factor in equation 2.

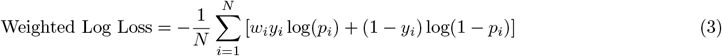

#### 4.5 Additional detail and data scope

SC2 enhances performance through model consensus and incorporates diverse KP to achieve superior outcomes. Unlike SC1, where models within each cluster share similar structures and the training data remains unchanged, SC2 introduces two distinct models, SC2-PB and SC2-PS, each utilizing different data, model structures, and data curation strategies. To ensure consistency, SC2 employs paired scalers to match feature spaces to their native scales. Specifically, SC2-PS intentionally excludes one feature to avoid performance degradation caused by potential loading mismatches. Although these two models use different input scales, they share a common output scale, enabling their results to be combined for consensus.

For SC2-PB, the model was trained on the PDBBind dataset and SC1 data to determine the optimal RMSD cutoff for performance evaluation, with an RMSD cutoff of 3 Å being found optimal in our case study (Table S4). Approximately 3.63 million data points were used for SC2-PB training, with an active-to-inactive imbalance ratio of 35.79. SC2-PB training data was curated with a higher proportion of decoys, improving the model’s ability to distinguish between ligands and decoys while extending the definition of actives beyond the conventional 2 Å threshold. The PDBScreen data for SC2-PS was curated differently, where only the most similar poses were retained from Glide SP docking data, rendering RMSD cutoff testing inapplicable. The specific data curation is detailed in the Equiscore[5] paper and in the SC2 portion of the PDBScreen database (519K),, where the imbalance ratio is approximately 6.19. In total, the SC2 data scope expanded to approximately 4.15M data points, a 55-fold increase over the SC1 75K dataset.

Both SC2-PB and SC2-PS shared the same Optuna parameter search settings, with optimal parameters determined based on test set performance. After evaluating various combinations, the two most promising models were finalized as SCORCH2.

#### 4.6 Inference speed

The SC2 framework requires atom-level feature computation, rendering its inference speed highly sensitive to receptor size. As a representative example, feature extraction was performed on the ADRB2 receptor (PDB ID: 3P0G), which contains only one single chain A. On a workstation equipped with an AMD Ryzen 9 7950X CPU utilizing 32 parallel processes, a throughput of approximately 100 receptor-ligand complexes per second was achieved. When the receptor was spatially truncated to a 12 Å binding pocket centered on the ligand (HET code: P0G), the feature extraction speed increased to approximately 180 complexes per second.

For model inference, performance was evaluated on the FNTA target, comprising approximately 53,000 data points and representing the largest compound cluster in the DUD-E dataset. On a server equipped with a single NVIDIA L40S GPU, inference using a single XGBoost model within SC2 required 0.29 seconds. When employing the consensus strategy—where predictions from both SC2-PS and SC2-PB models are integrated—the total inference time approximately doubled.

## Supporting information

SCORCH2 supplement material

## Supplementary information

Uploaded separately.

## Acknowledgements

For the purpose of open access, the author has applied a Creative Commons Attribution (CC BY) licence to any Author Accepted Manuscript version arising from this submission.

## Declarations

### Ethics approval and consent to participate Not applicable

### Data availability

Original PDBScreen data can be found at https://doi.org/10.5281/zenodo.8049380. PDBBind V2020 dataset could be found at https://www.pdbbind-plus.org.cn. BindingDB and Iridium dataset could be accessed from https://www.bindingdb.org/rwd/bind/index.jsp and https://www.eyesopen.com/iridium-database. DEKOIS, DUD-E could be found at https://uni-tuebingen.de/fakultaeten/mathematisch-naturwissenschaftliche-fakultaet/fachbereiche/pharmaziebiochemie/teilbereich-pharmazie-pharmazeutisches-institut/pharmazeutische-chemie/prof-dr-f-boeckler/dekois/ and https://dude.docking.org. Original VSDS-vd benchmark could be found at https://doi.org/10.5281/zenodo.13684010. Merck-FEP benchmark is commonly available at https://github.com/MCompChem/fep-benchmark. All evaluation data that supports this study can be found at https://zenodo.org/records/14994007.

### Code availability

The code used to reproduce this study is available under an MIT License via GitHub at https://github.com/LinCompbio/SCORCH2.

### Author contribution

L.C., D.H., and V.B. collaborated on the design of the study. L.C. spearheaded the method development and implementation, processed and collected training data, and conducted benchmark evaluations. P.B. provided additional review and practical insights. L.C., D.H., and V.B. drafted the paper, and all authors reviewed and approved the final version.

