## Supplementary material for "SCORCH2: a generalised heterogeneous consensus model for high-enrichment interaction-based virtual screening": SCORCH2 supplement material

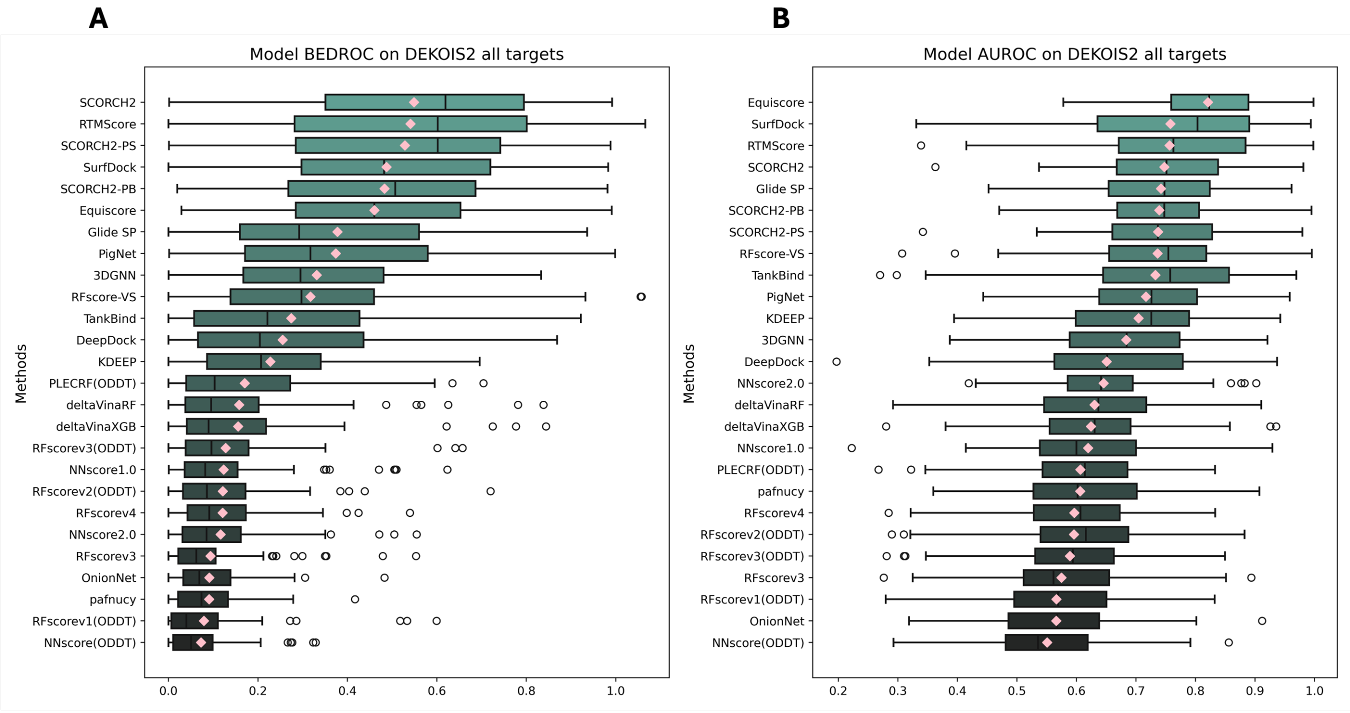


**Fig. S1**: Method AUROC (A) and BEDROC(B, $\alpha = 80.5$) on DEKOIS 2.0 dataset (data point n= 81). The pink diamonds denote the mean value for each metric, where a median line is included in each box-whisker plot. SCORCH2-PB - model trained with SC1 and PDBBind data, SCORCH2-PS - model trained with PDBScreen data, SC2 - model consensus by SCORCH2-PB and SCORCH2-PS.

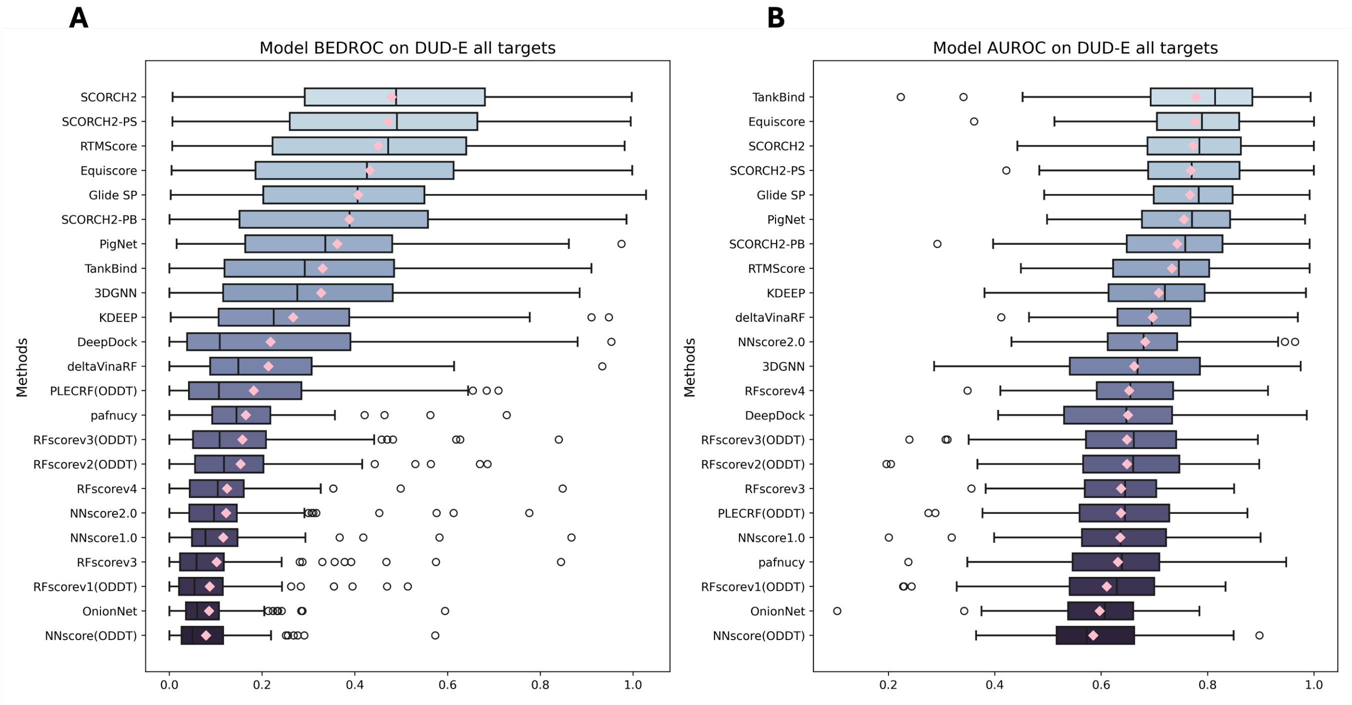


**Fig. S2**: Method AUROC (A) and BEDROC(B, $\alpha = 80.5$) on DUD-E dataset (data point n= 102). The pink diamonds denote the mean value for each metric, where a median line is included in each box-whisker plot. SCORCH2-PB - model trained with SC1 and PDBBind data, SCORCH2-PS - model trained with PDBScreen data, SC2 - model consensus by SCORCH2-PB and SCORCH2-PS.

**Removed PDBIDs from PDBScreen**

| | | | | | | | | | |
|---|---|---|---|---|---|---|---|---|---|
| 1d3g | 1li4 | 1n4h | 1nq7 | 1nrl | 1pwl | 1r9o | 1t40 | 1xu9 | 1z3n |
| 1z89 | 1z8a | 2dux | 2duz | 2fpt | 2fqi | 2fvj | 2fz8 | 2fz9 | 2g6i |
| 2g6j | 2gtk | 2hc4 | 2hv5 | 2hvn | 2i16 | 2i17 | 2ikh | 2iki | 2ipw |
| 2is7 | 2npa | 2p54 | 2pd5 | 2pdb | 2pdf | 2pdi | 2pdj | 2pdk | 2pdm |
| 2pdp | 2pdu | 2pdx | 2pev | 2pfh | 2prg | 2prl | 2pzn | 2r24 | 2rbe |
| 3bzu | 3ch6 | 3cwd | 3czr | 3dct | 3dn5 | 3et3 | 3fei | 3fej | 3frj |
| 3fur | 3g0u | 3g5e | 3g8i | 3g9e | 3ghr | 3ghs | 3ght | 3ghu | 3gyt |
| 3gyu | 3h6k | 3hfg | 3hvl | 3ipq | 3ips | 3ipu | 3jwu | 3jwv | 3jx0 |
| 3jx3 | 3jx5 | 3kvk | 3kvm | 3lbo | 3lep | 3lmp | 3lqg | 3lql | 3lz3 |
| 3lz5 | 3m0i | 3m4h | 3m64 | 3mb9 | 3mc5 | 3n5v | 3n5w | 3n5x | 3n5y |
| 3n60 | 3n61 | 3n62 | 3n63 | 3n64 | 3n65 | 3n66 | 3nlj | 3nln | 3nlq |
| 3nlz | 3nny | 3okh | 3oki | 3olf | 3omk | 3omm | 3onc | 3oof | 3ook |
| 3p89 | 3png | 3qt0 | 3rqk | 3rqm | 3rx2 | 3rx3 | 3rx4 | 3svp | 3t03 |
| 3t42 | 3tyo | 3u2c | 3ufr | 3ufv | 3zwt | 4c7j | 4cao | 4cdt | 4ctr |
| 4ctw | 4ctx | 4cx4 | 4cx5 | 4dk7 | 4dk8 | 4dm6 | 4dm8 | 4eux | 4gca |
| 4gq0 | 4igs | 4iju | 4ijv | 4ims | 4imw | 4js3 | 4jsf | 4jsh | 4jsj |
| 4jts | 4jtt | 4jtu | 4jyg | 4jyi | 4k1l | 4k5d | 4k5e | 4kch | 4lau |
| 4laz | 4lb3 | 4lb4 | 4lbr | 4lbs | 4ls2 | 4nkc | 4nz2 | 4oqv | 4prt |
| 4puu | 4q7b | 4qbx | 4qr6 | 4qx4 | 4qxi | 4rpq | 4ruj | 4uda | 4udb |
| 4ugz | 4uh2 | 4uh3 | 4upm | 4v3w | 4y29 | 4ys1 | 4yu1 | 4zmg | 5a86 |
| 5ad4 | 5ad5 | 5ad7 | 5ad8 | 5adb | 5ade | 5avi | 5dwl | 5fvq | 5g0n |
| 5h73 | 5ha7 | 5k7k | 5k9c | 5k9d | 5l7e | 5l7g | 5mvc | 5nky | 5ow7 |
| 5ow9 | 5owd | 5pgu | 5pgw | 5pgy | 5q0j | 5q0l | 5q0m | 5q0n | 5q0o |
| 5q0p | 5q0q | 5q0r | 5q0s | 5q0t | 5q0v | 5q0w | 5q0x | 5q0y | 5q0z |
| 5q10 | 5q11 | 5q12 | 5q13 | 5q14 | 5q15 | 5q16 | 5q18 | 5q19 | 5q1a |
| 5q1b | 5q1c | 5q1d | 5q1e | 5q1f | 5q1g | 5q1h | 5q1i | 5qii | 5unt |
| 5unu | 5unv | 5unw | 5uo0 | 5vui | 5vuj | 5vun | 5vup | 5vur | 5vus |
| 5vut | 5vuu | 5w0c | 5w49 | 5x23 | 5x24 | 5xxi | 5y44 | 5ycp | 5yp6 |
| 5zf8 | 6aur | 6cjf | 6e3g | 6et4 | 6fo7 | 6fo8 | 6fo9 | 6fob | 6fod |
| 6gg8 | 6hty | 6ijs | 6ilq | 6jme | 6lp6 | 6lp7 | 6pn0 | 6pn1 | 6pn4 |
| 6pn7 | 6pn8 | 6ssq | 6syw | 6t3p | 6tuf | 6w9h | 6w9i | 6xp9 | 7bpy |
| 7bpz | 7bq0 | 7bq2 | 7bq3 | 7bq4 | 7kxd | 7kxf | | | |
| Total: 317 | | | | | | | | | |

**Table S1**: List of PDBIDs removed from SC2-PS training data that duplicate with DEKOIS 2.0 and DUD-E unseen targets.

| Target Name | PDB ID | UniProt ID |
|---|---|---|
| ALDR | 2HV5 | P15121 |
| CP2C9 | 1R9O | P11712 |
| CP3A4 | 3NXU | P08684 |
| DHI1 | 3FRJ | P28845 |
| HXK4 | 3F9M | P35557 |
| KITH | 2B8T | Q9PPP5 |
| NOS1 | 1QW6 | P29476 |
| PGH1 | 2OYU | P05979 |
| PPARA | 2P54 | Q15788 |
| PPARG | 2GTK | Q15788 |
| PYRD | 1D3G | Q02127 |
| SAHH | 1LI4 | P23526 |
| **SC2-PB duplicated data** | | |
| CP2C9 | 5K7K | P11712 |
| SAHH | 3NJ4 | P23526 |
| DHI1 | 3CH6 | P28845 |
| NOS1 | 4KCL | P29476 |

**Table S2**: DUD-E targets that do not appear in the PDBBind V2020 dataset, and proteins with the same UniProt ID in the SC2-PB training data.

| Target Name | PDB ID | UniProt ID |
|---|---|---|
| 11BETAHSD1 | 3TFQ | P28845 |
| ACE2 | 1R4L | Q9BYF1 |
| ALR2 | 1AH3 | P80276 |
| COX1 | 3KK6 | P05979 |
| CYP2A6 | 1Z11 | P11509 |
| ER-BETA | 3OLL | Q15788 |
| INHA | 1P44 | P9WGR1 |
| MMP2 | 1HOV | P08253 |
| PPARA | 2P54 | Q15788 |
| PPARG | 1FM9 | Q15788 |
| TK | 1W4R | P04183 |
| **SC2-PB duplicated data** | | |
| CYP2A6 | 2FDW | P11509 |
| 11BETAHSD1 | 3CH6 | P28845 |

**Table S3**: DEKOIS 2.0 targets that do not appear in the PDBBind B2020 dataset, and proteins with the same UniProt ID in the SC2-PB training data.

**Table S4**: Performance comparison between different model combinations based on the DEKOIS 2.0 and DUDE benchmarks, Glide SP docking poses. The table reports the AUC-ROC, BEDROC ($\alpha$=80.5), and EF (Enrichment Factor) for each dataset, the best combination is bolded.

| Model | AUC-ROC | BEDROC (80.5) | EF 0.5% | EF 1% | AUC-ROC | BEDROC (80.5) | EF 0.5% | EF 1% |
|---|---|---|---|---|---|---|---|---|
| | DEKOIS | | | | DUDE | | | |
| SC2-PS, rmsd cutoff 2 | 0.7371 | 0.5287 | 21.0957 | 18.6206 | 0.7684 | 0.4717 | 23.2832 | 19.5371 |
| SC2-PB, rmsd cutoff 2 + SC2-PS | 0.7402 | 0.5382 | 20.6965 | 19.1030 | 0.7688 | 0.4714 | 23.2030 | 19.6013 |
| SC2-PB, rmsd cutoff 2.5 + SC2-PS | 0.7426 | 0.5379 | 21.3426 | 19.0891 | 0.7701 | 0.4727 | 23.3672 | 19.6597 |
| SCORCH2 | **0.7477** | **0.5486** | **21.6220** | **19.2636** | **0.7728** | **0.4791** | **23.5229** | **19.9283** |

**Table S5**: XGBoost Hyperparameters Used in Model Training

| Parameter | Value/Range | Description |
|---|---|---|
| `tree_method` | `hist` | Algorithm used to construct trees. |
| `device` | `cuda` | Device for training. |
| $\lambda$ | log-uniform(0.001, 10.0) | L2 regularization term (Ridge regression). |
| $\alpha$ | log-uniform(0.001, 10.0) | L1 regularization term (Lasso regression). |
| `colsample_bytree` | uniform(0.1, 1.0) | Fraction of features to sample for each tree. |
| `subsample` | uniform(0.3, 1.0) | Subsample ratio of the training instances. |
| `max_depth` | integer(1, 30) | Maximum depth of a tree. |
| `min_child_weight` | integer(1, 30) | Minimum sum of instance weight (hessian) needed in a child. |
| `n_estimators` | integer(1, 3000) | Number of gradient boosted trees. |
| `learning_rate` | uniform(0.0001, 0.1) | Learning rate pool. |
| `scale_pos_weight` | calculated | Balancing of positive and negative weights. |
| `objective` | `binary:logistic` | Learning task and objective function. |

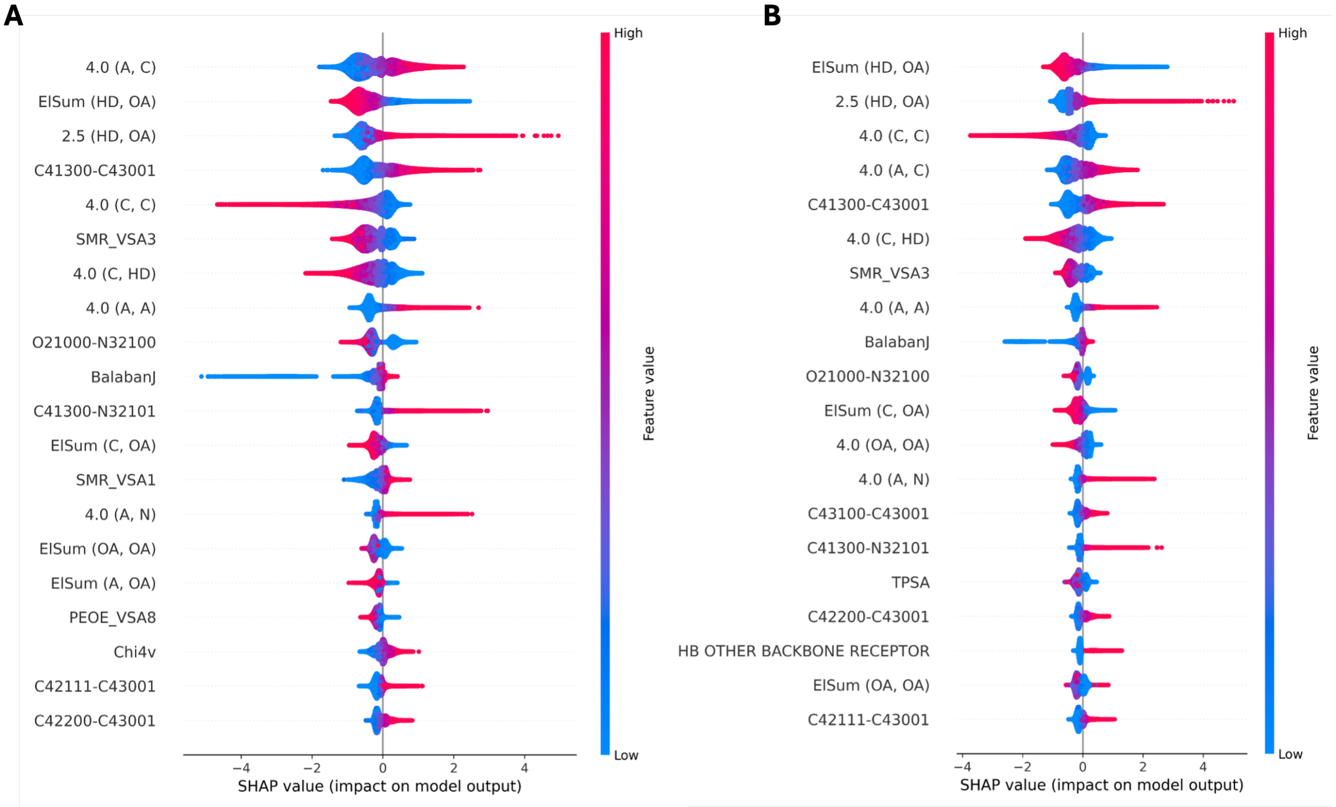


**Fig. S3**: SHAP value beeswarm plot visualization. A, SC2-PB, rmsd cutoff - 2.5 Å; B, SC2-PB, rmsd cutoff - 2 Å. The color indicates the value of each feature (blue for lower numbers and red for higher numbers), while the thickness of the line corresponds to the number of samples. Features are ranked by importance, with the most important feature at the top.

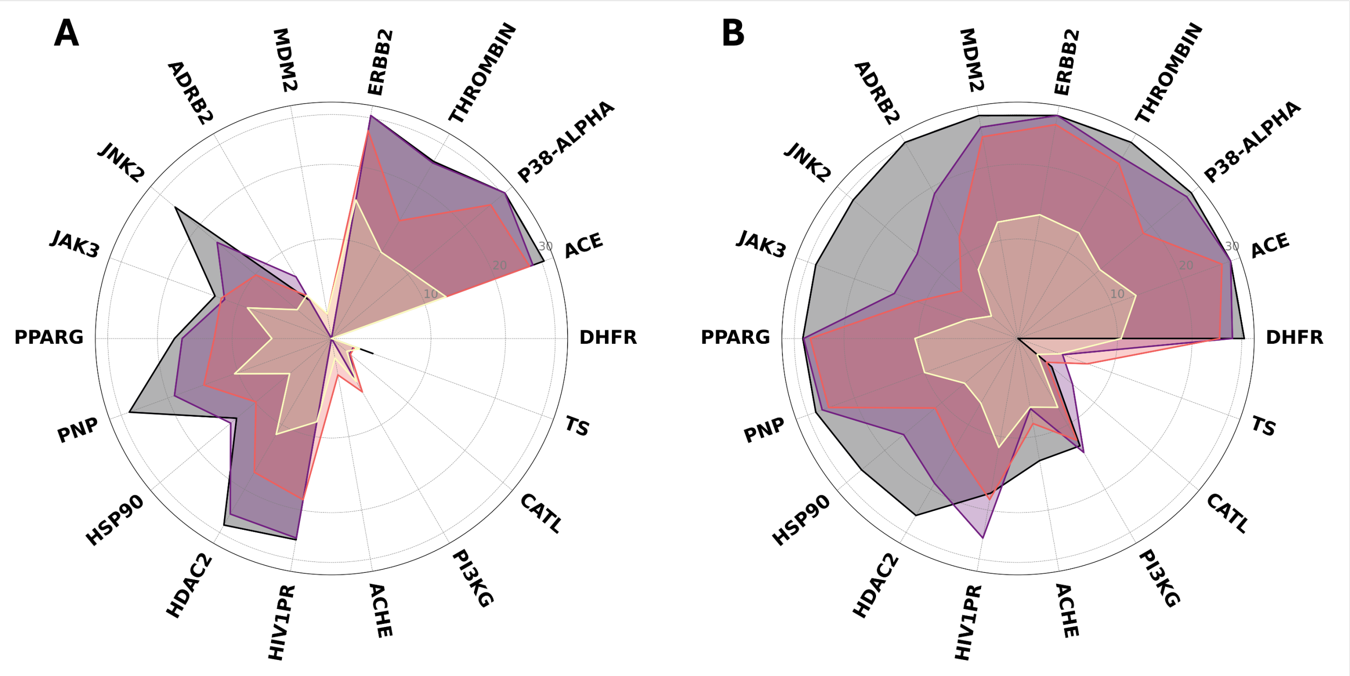


**Fig. S4**: Screening performance of SC1 (A) and SC2 (B) on the subset of DEKOIS 2.0 (data point n=18). The label on the outer circle indicates the target name. Within each circle, the charcoal grey line, violet line, reddish-orange line, and pale gold line represent EF values at 0.5%, 1%, 2%, and 5%, respectively. The scale at the top of each circle indicates the absolute enrichment value.

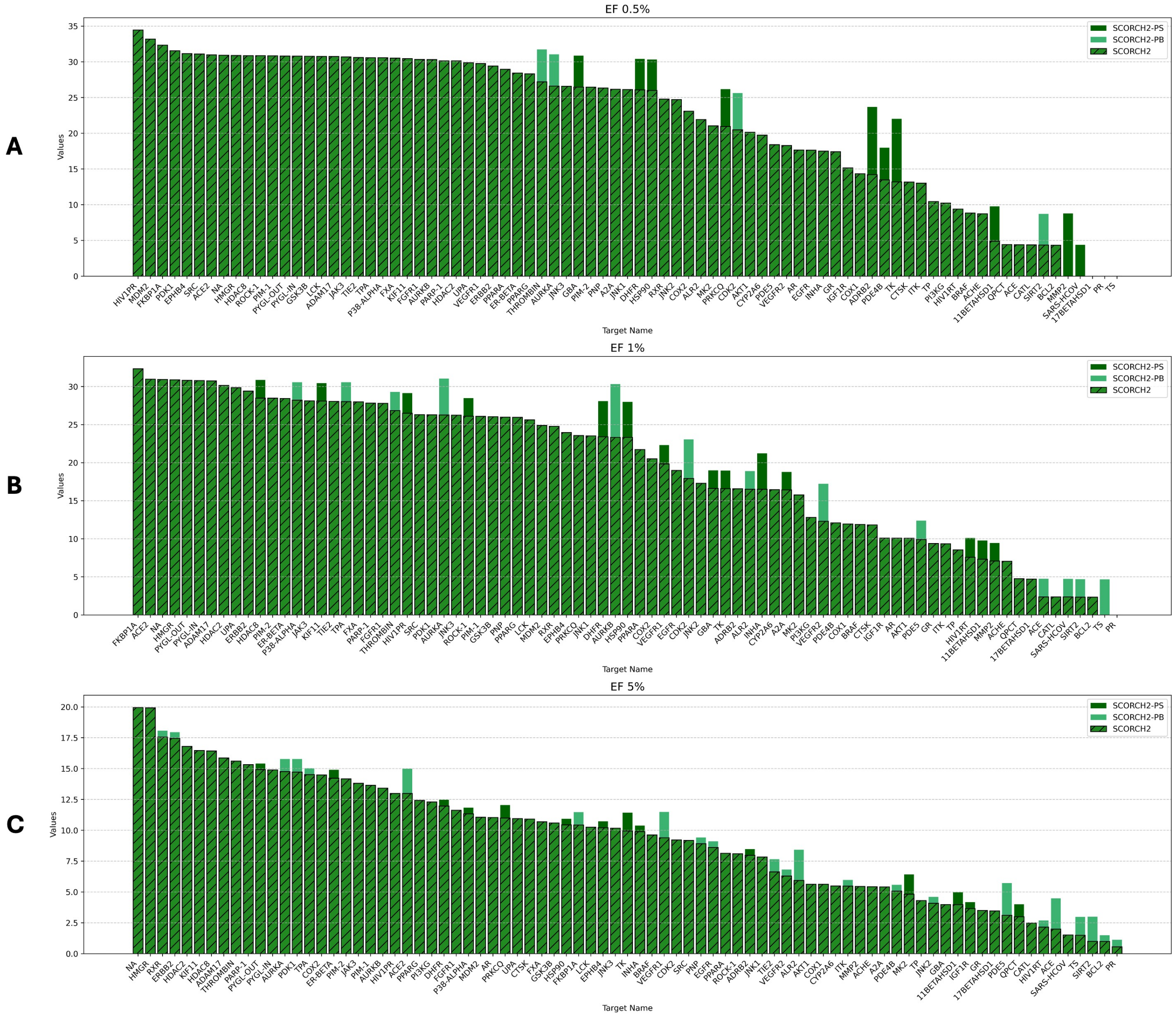


**Fig. S5**: Stacked bar plot of SC2 VS performance at target level. The x-axis represents the names of all targets in the DEKOIS 2.0 dataset (data point n=81), while the y-axis denotes the absolute number of EF. A - EF at top 0.5% level; B - EF at - top 1% level ; C - EF at - top 5% level. And SC2-PS single VS result is represented in dark green, same as the mediumseagreen for SC2-PB result, and SC2 (model consensus) result is represented with bars in forest green with dark hatch.

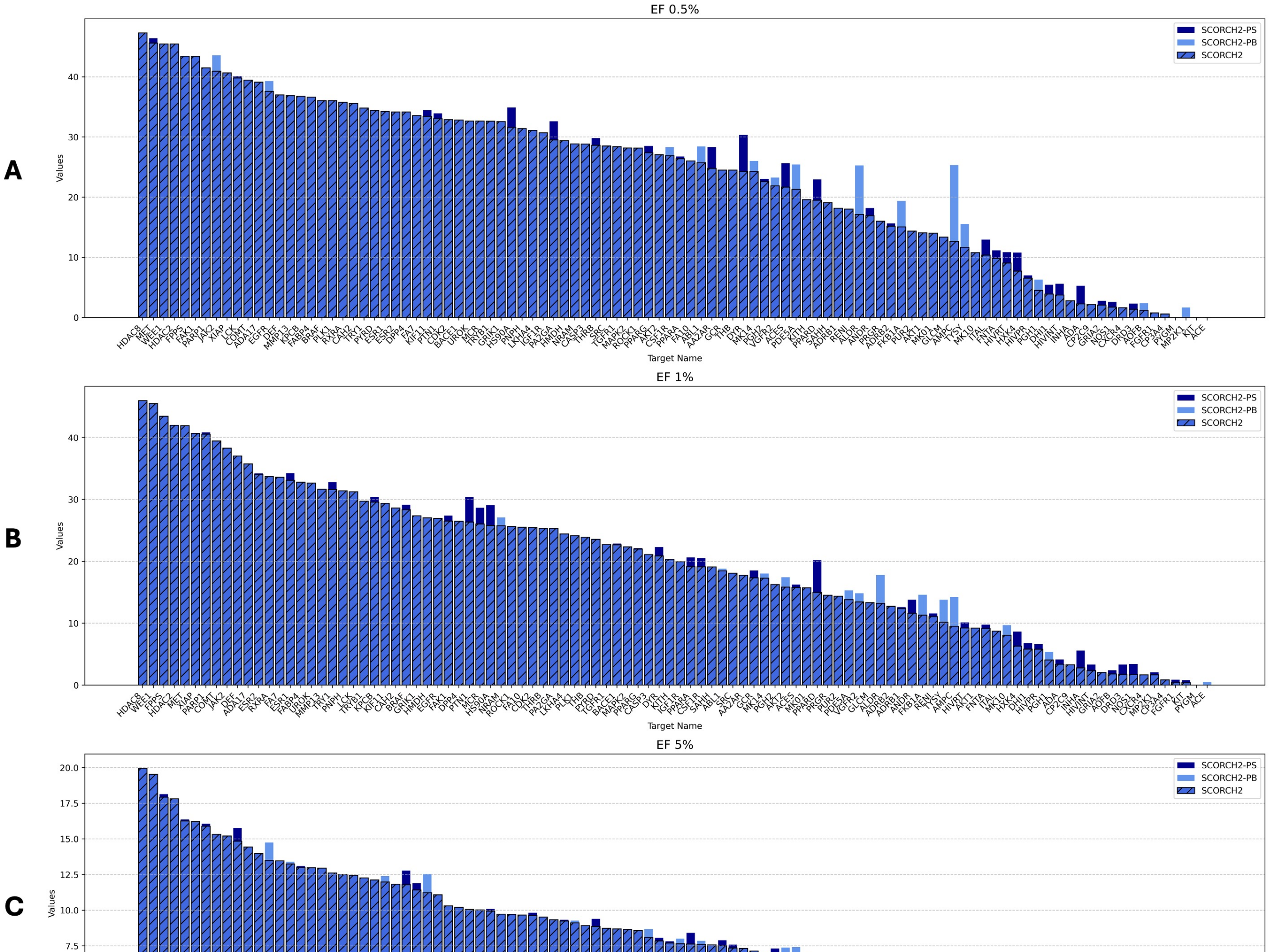


**Fig. S6**: Stacked bar plot of SC2 VS performance at target level. The x-axis represents the names of all targets in the DUD-E (data point n=102), while the y-axis denotes the absolute number of EF. A - EF at top 0.5% level; B - EF at - top 1% level; C - EF at - top 5% level. And SC2-PS single VS result is represented in dark blue bar, same as the cornflower blue bar for SC2-PB result, and SC2 (model consensus) result is represented with royal blue bars with dark hatch.

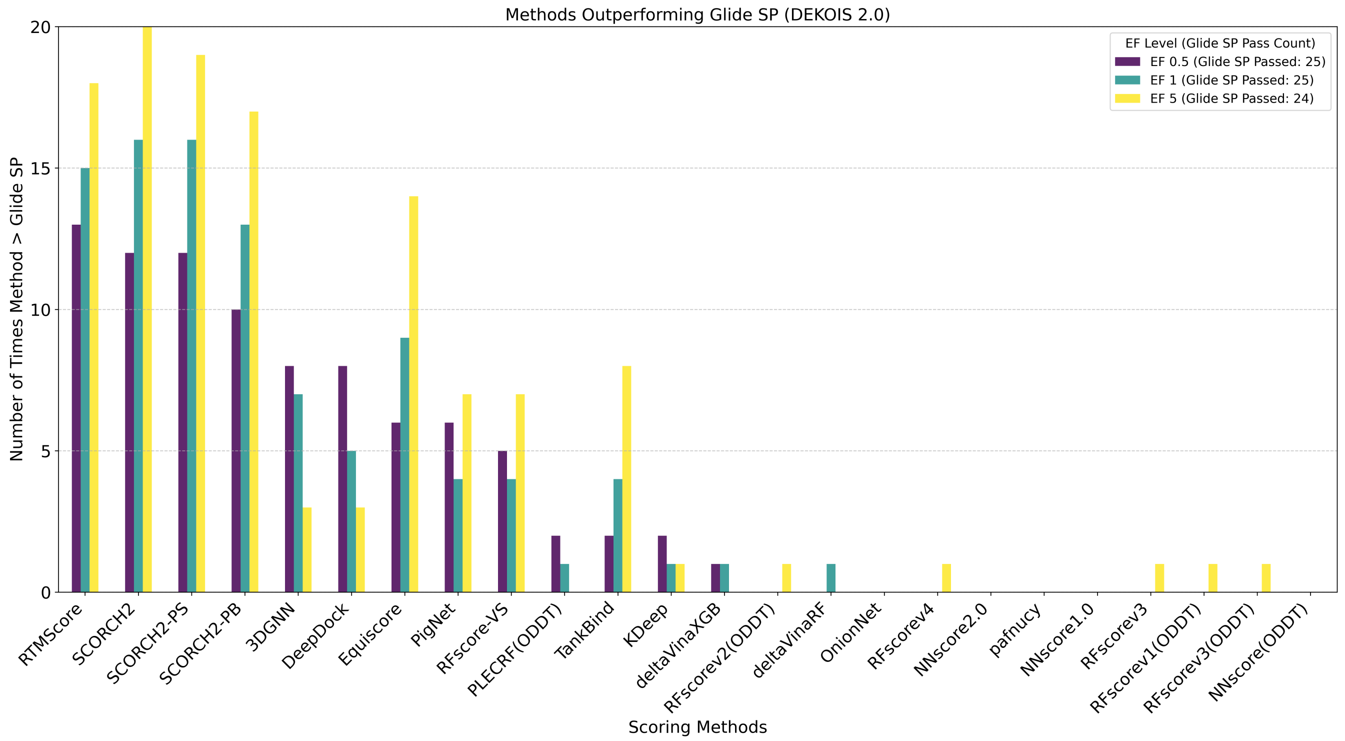


**Fig. S7**: Bar plot illustrating the marginal performance improvement on the DEKOIS 2.0 dataset. The x-axis represents the names of all scoring methods, while the y-axis denotes the absolute number of enrichment factor (EF) occurrences at different thresholds: (A) EF at the top 0.5% level ($n = 25$); (B) EF at the top 1% level ($n = 25$); (C) EF at the top 5% level ($n = 24$). Purple, dark green, and yellow bars indicate the number of targets for which each method outperforms the native Glide score.

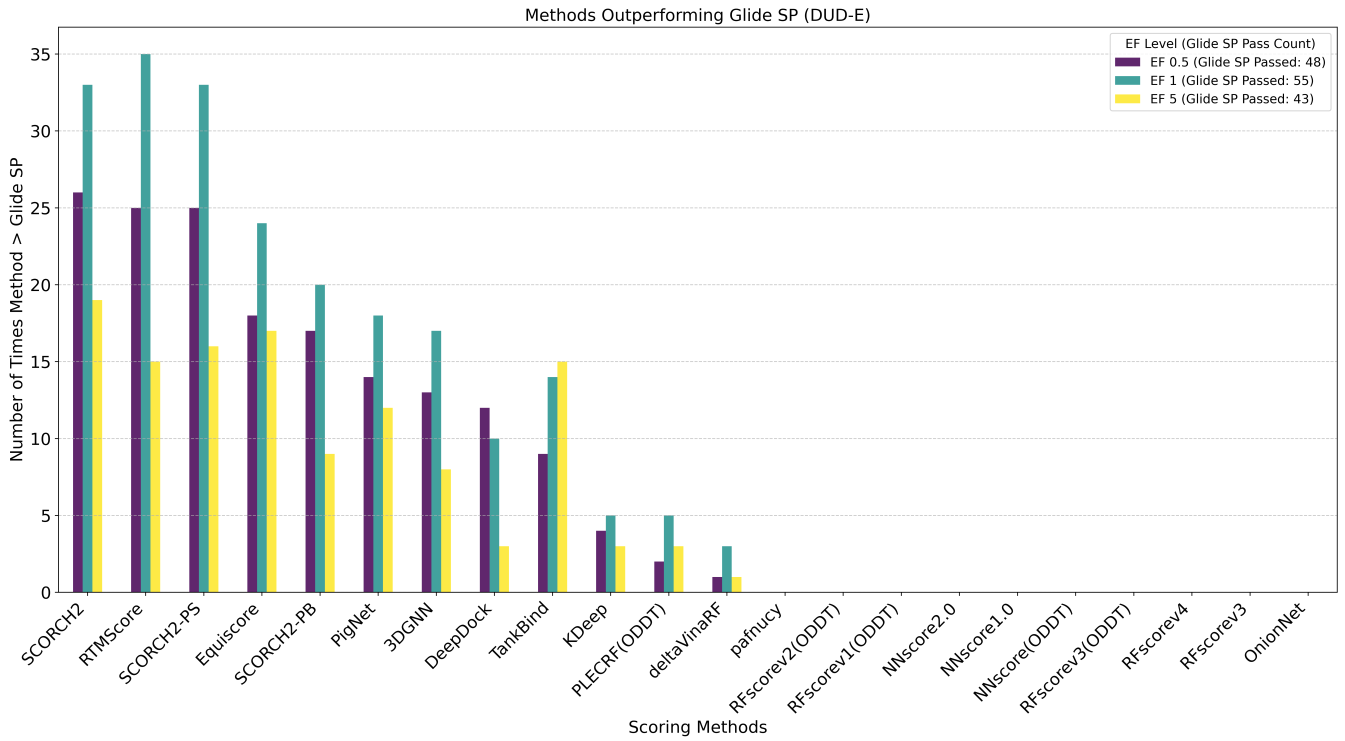


**Fig. S8**: Bar plot illustrating the marginal performance improvement on the DUD-E dataset. The x-axis represents the names of all scoring methods, while the y-axis denotes the absolute number of enrichment factor (EF) occurrences at different thresholds: (A) EF at the top 0.5% level ($n = 48$); (B) EF at the top 1% level ($n = 55$); (C) EF at the top 5% level ($n = 43$). Purple, dark green, and yellow bars indicate the number of targets for which each method outperforms the native Glide score.
